# Molecular dynamics insights into biomineralisation mediated by acidic intrinsically disordered proteins: a case study of molluscan Aspein from the pearl oyster *Pinctada fucata*

**DOI:** 10.64898/2026.06.03.729799

**Authors:** Julien Mignon, Antonio Monari, Sonia Longhi, Catherine Michaux

## Abstract

Biomineralisation is a ubiquitous phenomenon that still fascinates the scientific community. Mineralised structures are most notably encountered in marine lifeforms the shell or exoskeleton of which are formed by precipitating specific calcium carbonate (CaCO_3_) polymorphs, known as amorphous (ACC), vaterite, aragonite, and calcite. To control crystalline polymorphism in their shell layers, bivalves have evolved strategies involving ion-binding secretomes composed of shell matrix proteins (SMPs). Secreted in the prismatic layer of the pearl oyster *Pinctada fucata*, Aspein is an unusually acid-rich protein exclusively aspartic and famously recognised as the most acidic SMP known to date. Through *in vitro* crystallisation experiments, Aspein was shown to select calcite over aragonite in modern seawater conditions. However, how it exerts its selectivity with respect to its structural properties remains enigmatic. By combining sequence-based predictions with all-atom molecular dynamics simulations, we unveiled that Aspein is an intrinsically disordered organic matrix protein (IDOMP) undergoing ion-specific chain collapse and phase separation. Thanks to its unique Asp density and conformational fuzziness, Aspein first sequesters Ca^2+^ then attracts CO_3_^2−^ ions before stabilising ACC and calcite-like phases, while precluding the incorporation of Mg^2+^ to avoid aragonite formation. Indeed, via Asp-Ca^2+^ bridges, structure-less clusters coalescence into supersaturated protein-CaCO_3_ assemblies initiates calcite crystallisation. Therefore, Aspein plasticity, condensation susceptibility, and aspartic density critically modulate the pathway that defines CaCO_3_ polymorphism by inhibiting nucleation and controlling the growth of biogenic minerals. From a broader perspective, our study underpins some of the molecular and physico-chemical principles governing biomineralisation processes mediated by IDOMPs, the overlooked, yet prevalent, role of which was mechanistically unexplored.

## 1. Introduction

Acidic proteins, characterised by an enrichment in either aspartate (Asp), glutamate (Glu), or both amino acids are predominant regulators of nucleic acid recognition and of the physiological metallic cations homeostasis. These functions are ubiquitously encountered across all the domains of life.^1–3^ Amongst such ubiquitous processes, biomineralisation takes advantage of the unique binding, sequestering, and clustering abilities of acidic proteins. Consisting in the compartmented intra- and/or extracellular formation and deposition of inorganic minerals by living organisms, biomineralisation is a highly regulated process that has independently occurred multiple times over the course of evolution in pro- and eukaryotes.^4,5^ In marine environments, tissues commonly mineralise through the precipitation of calcium carbonate (CaCO_3_).^6,7^ With respect to the ordered ionic arrangement adopted within the crystal unit cell, CaCO_3_ can nucleate and grow into three distinct polymorphs, from the least to the most stable: vaterite, aragonite, and calcite.^8^ Acting as precursors and kinetic modulators of biogenic crystalline forms, CaCO_3_ can also be stabilised into amorphous phases, accordingly referred to as amorphous calcium carbonate (ACC).^9,10^

Such structural and morphological diversity is observed in bacteria, but most noteworthily in crustaceans and scleractinian corals, as well as brachiopods and molluscs which exploit it for the generation of their exoskeleton and shell, respectively. Indeed, with the help of Ca^2+^-binding proteins and other organic polymers, they are able to select distinct polymorphs and crystal morphologies according to the targeted mineralised tissues.^11–15^ Since completion of its genome, one of the most established model organisms for studying marine biomineralisation processes is the Japanese pearl oyster *Pinctada fucata* (*P*. *fucata*).^16,17^ As for other members of the Mollusca phylum, the shell of *P*. *fucata* is organised into three mineralised structures that are all constituted from specific CaCO_3_ polymorphs. The inner zone of the mantle is characterised by an aragonitic nacreous layer, while the prismatic layer located at the outer edge of the mantle is made of calcite prisms.^18,19^ Additionally, the valve hinge ligament is composed of fibrous and needle-shaped aragonite crystals.^20^

While molluscan shells are prevalently inorganic (∼95% CaCO_3_), polymorph selection and other aspects of their formation are notably regulated by shell matrix proteins (SMPs). Expressed by specialised epithelia in the mantle before diffusing to the extrapallial space where they self-assemble with inorganic materials and other organic molecules (polysaccharides and lipids), they create an intricate extracellular organic matrix where biomineralisation can occur.^7,21–23^ Illustratively, 72 of such organic matrix proteins have been identified in the shell layers of *P*. *fucata*.^24^ Amongst SMPs, those being acid soluble and containing a high level of Asp/Glu amino acids are unique agents that finely modulate the adsorption, nucleation, crystallisation, inhibition, morphology, and deposition of CaCO_3_.^25–27^ In such regulatory purposes, prismatic and nacreous layers have developed diversified and dedicated acidic secretory repertoires with respect to seawater conditions.^28^ Current oceans have indeed a high Mg^2+^/Ca^2+^ ratio of ∼5.0−5.2, qualifying seawater chemistry as aragonitic given that, under the influence of Mg^2+^ cations, stochastic nucleation and precipitation of the aragonite polymorph is thermodynamically favoured at such ionic concentrations.^29–31^ As both the nacre and prisms are deposited from the extrapallial fluid subjected to such high Mg^2+^ levels, acid-rich prismatic SMPs hence help capturing Ca^2+^ for calcite crystallisation.

Historically discovered by Tsukamoto D., *et al*. in 2004, the *aspein* gene transcript encodes an unusually acidic SMP exclusively secreted by the epithelial tissues of the outer mantle edge corresponding to the prismatic layer of *P. fucata*.^32^ Amongst SMPs, Aspein is peculiar owing to its exceptionally low isoeletric point (pI∼1.45) arising from its aspartic density, making it the most acidic SMP known to date.^21^ Sequence-wise, Aspein is indeed characterised by a remarkable acidic content intriguingly biased towards Asp amino acids, as it possesses 238 (60.4%) of Asp for only 6 (1.5%) Glu. In addition, high serine (Ser) and glycine (Gly) levels account for 13.2 and 16.0% of the total amino acid content, respectively. Such an extreme sequence composition classifies Aspein as an unusual acid-rich protein (ARP), comparable to Asp-rich proteins (Asprich).^33^ Unsurprisingly, the Asprich family encompasses several marine SMPs and functionally diversifies into metal regulation, and, most meaningfully, biomineralisation.^34^

Through *in vitro* crystallisation assays conducted with a truncated version (Aspein-D1) under the physico-chemical Mg^2+^/Ca^2+^ parameters met in the extrapallial space, Aspein was later revealed by Takeuchi T., *et al*. to modulate calcite nucleation, precipitation, and morphology by stabilising ACC and selectively binding Ca^2+^ cations over Mg^2+^ via its crucial polyAsp (polyD) domain.^35^ Based on their results and the shell layer secretion of Aspein, the acidic SMP was propounded to act as an accumulator of Ca^2+^ and CO_3_^2−^ ions in a disordered prenucleation state (i.e. ACC) by diffusing throughout the extrapallial fluid before aggregating inside the prismatic wall where it accelerates ACC transformation into calcite. Given its polyelectrolytic nature, Aspein is expected and was suggested to be intrinsically disordered, although no compelling *in vitro* nor *in silico* evidence has been provided hitherto.^35,36^ Intrinsically disordered proteins (IDPs), existing as a dynamical ensemble of interconverting conformers, are highly relevant to biomineralisation, as emphasised by their plasticity facilitating ions clustering. An increasing proportion of SMPs have indeed been recognised to exhibit characteristics of unstructured proteins.^17,36–42^ Liquid-liquid phase separation (LLPS) phenomena, chiefly driven by interactions engaging intrinsically disordered (IDRs) and low complexity regions (LCRs), are now more than ever emerging as a new paradigm for apprehending the generation of biologically active proteinaceous assemblies; and biomineralisation is no exception. Yet only recently, studies have begun to pinpoint the interplay between SMPs and ionic species into heterotypic protein-inorganic condensates for regulating the crystallisation of biominerals.^40,43–45^ Notwithstanding such insights, the literature on Aspein, and more generally disordered SMPs, that we propose here to designate under the term "*intrinsically disordered organic matrix protein (IDOMP)*", is very scarce since their discovery. This is particularly true concerning structural data and the pathways through which they self-assemble with minerals within organic frameworks remain highly elusive. In this respect, biomineralisation processes are still mechanistically enigmatic, and this deepens when introducing IDP into the picture. While little attention is gradually given to the role of intrinsic disorder and LLPS in protein-mediated biomineralisation, IDOMPs remain largely overlooked in recent reviews and studies due to the challenges raised by their structural heterogeneity. As such, the molecular pathways associated with their sequence and conformational properties related to the precipitation of biogenic minerals are also very much unexplored by experimental and/or computational meanings.

Altogether, these observations prompted us to unravel the principles driving calcite biomineralisation, using Aspein from the pearl oyster *P*. *fucata* as a model acid-rich (Asprich) IDOMP. Through the combination of sequence-based prediction with all-atom equilibrium molecular dynamics (MD), we highlighted unique features of the Aspein protein related to intrinsic disorder and phase separation in ionic conditions mimicking the Mg^2+^/Ca^2+^ imbalance encountered within modern seawater and the extrapallial fluid of marine bivalves. Comparing the findings from our computational simulations with the experimental works published so far, the present study aims at shedding new light on the molecular basis governing CaCO_3_ biomineralisation in marine lifeforms and, most especially, on the regulatory role of acidic IDOMPs in polymorph selection for molluscan shell formation.

## 2. Methods

### 2.1. Bioinformatics and sequence-based prediction

All properties were determined or predicted from the mature sequence of full-length wild-type Aspein secreted by *P*. *fucata* and indexed in the UniProt database under the Q76K52 entry. Sequence composition and isotopically averaged molecular mass (M) were provided by the Protein Tool from the Prot pi webserver. Disorder scores were calculated along the query sequence by exploiting several IDP-oriented predictors: DisoMine, DisoPred3, metapredict (v3.0), PrDOS, RFPR-IDP, as well as RIDAO, encompassing PONDR and IUPred tools.^46–52^ Predicted per-residue percentage of intrinsic disorder (PPID) was averaged over the predictors. Complementarily, cumulative distribution function (CDF) and charge-hydropathy (CH) plots were extracted from the PONDR platform.^53^ Charge-driven conformational preferences were assessed by localising the protein on the Das-Pappu state diagram, generated via the Classification of Intrinsically Disordered Ensemble Regions (CIDER) webserver.^54^ Linear distribution of the net charge per residue and hydropathy was also computed and plotted over a sliding window of five amino acids with CIDER. Occurrence and localisation of LCRs and molecular recognition features (MoRFs) were assessed with the PlaToLoCo and MoRFchibi tools, respectively.^55,56^ Secondary structure propensities were determined with PSIPRED and NetSurfP3.0, whereas phosphosites were predicted from the DEPP (DisPhos) and NetPhos3.1a servers.^47,57–59^ Regarding LLPS-oriented and interaction properties, we concatenated the predicted scores and profiles generated with the following algorithms: deePhase, FuzDrop, FuzPred, MolPhase, and ParSe.^60–64^

### 2.2. Preparation of atomistic systems prior to simulation

The sequence of full-length wild-type Aspein from *P*. *fucata* was retrieved from the UniProt database and stripped from its signal peptide (residue 1 to 19) and part of its polyD domain (residue 131 to 413) to match the truncated version experimentally assessed by Takeuchi T., *et al*.^35^ From this 111 amino acids long sequence (Aspein-D1), the initial tertiary structure of Aspein was modelled with the combined MMseqs2-AlphaFold2 approach freely available on the ColabFold platform.^65^ Amongst the top ranked models, the structure with the less biased intramolecular contacts was selected for subsequent file preparation. By exploiting the CHARMM-GUI suite, sidechain atoms were properly protonated, and termini were capped with the PDB Reader & Manipulator tool.^66,67^ The protein chain was centred in a cubic water box with a 10 Å buffer of water molecules added to its edges. Three ionic environments were considered: i) 10 mM NaCl, ii) 10 mM CaCO_3_, and iii) 10 mM CaCO_3_ with 50 mM MgCl_2_ **(Figure S1)**, the latter corresponding to the concentrations and Mg/Ca ratio of the extrapallial fluid. For each system, the required number of ionic species was added with respect to the box volume and protein net charge. To assess phase separation properties, SG(D)3 and SG(D)4 stretches from the polyD domain of Aspein were generated in an extended random coil conformation before being capped and protonated using the Builder plugin in PyMOL.^68^ Within 100 Å-sided water cubic boxes, 20 copies of each segment, for a total of 40 peptides, were randomly dispersed at a minimal 10 Å distance apart from each other, as well as solvated and neutralised in i) 50 mM NaCl, ii) 50 mM CaCO_3_, and iii) 50 mM CaCO_3_ with 250 mM MgCl_2_ **(Figure S1)** using CHARMM-GUI Multicomponent Assembler.^69^ All system parameters are summarised in Table S1.

### 2.3. Simulation of protein-ions systems by all-atom equilibrium molecular dynamics

While acknowledging the limitations of current non-polarisable force fields with regard to the parameters of divalent cations, we opted for a compromise between the description of the protein chain and ionic properties. Drawing inspiration from previous biomineralisation-oriented MD studies and with respect to its reliability to capture the conformational dynamics of IDPs in their environmental conditions, the CHARMM36m force field was therefore selected for modelling the protein atoms and ions (Na^+^, Ca^2+^, Mg^2+^, Cl^−^, and CO_3_^2−^) in conjunction with the TIP3P water model modified for CHARMM.^70–73^ MD simulations were conducted in independent triplicates with the GROMACS 2023.1 suite implemented with GPU (CUDA 11.7.0) accelerators.^74,75^ Newton’s equations of motion were integrated with a 4 fs timestep by applying hydrogen mass repartition (HMR) on the initial topologies, allowing to reduce high-frequency motion.^76,77^ In addition, the LINCS algorithm was used to constrain bonds involving hydrogen atoms to their equilibrium length.^78^ Each system was first minimised for a maximum of 5000 steps using the steepest descent algorithm before being thermalised, equilibrated, and propagated in the isothermal and isobaric (NPT) ensemble with the leap-frog integrator **(Figure S1)**.^79,80^ Pressure of 1 bar and temperature of 300 K were enforced by a C-rescale barostat and a modified Berendsen thermostat (V-rescale), respectively.^81,82^ Short-range non-bonded interactions were computed with a distance cut-off of 1.2 nm, whereas Particle Mesh Ewald (PME) summation was used for long-range coulombic interactions.^83^ Production MD replicates were propagated in periodic boundary conditions (PBC) in all three dimensions for a duration of 1 µs each. Atom coordinates and energies were recorded every 100 ps along the trajectories.

Modules directly invoked from the GROMACS suite were conjunctively used with in-house built Python scripts for analysis and clustering. For each system, time-dependent and per-residue variables, and their associated probability density functions (PDF) were computed and averaged across the replicates. Root-mean square deviation (RMSD) was calculated from the equilibrated structure, while root-mean square fluctuation (RMSF) was determined for each amino acid by taking the time-averaged structure as reference. Over the replicates, a total of 18000 frames were clustered by defining a 5.0 Å RMSD cut-off between the nearest neighbours. Principal component analysis (PCA) was carried out using the RMSD of backbone atoms for building the covariance matrix. Along the two first principal components capturing the essential motions of sampled systems (PC1 and 2), probabilities were computed and distributed in a hundred of basins with the kernel density estimation. Regarding secondary structure classes, they were defined using the STRIDE algorithm implemented in the Timeline VMD plugin: turns (i-i+3 turn), extended β-sheet, isolated β-bridge (usually i-i+>6), α-helix (i-i+4 helix), 310-helix (i-i+3 helix), π-helix (distorted or bulged i-i+5 helix), and coil. Trajectory visualisation and snapshot rendering were performed with the VMD molecular graphic program.^84^

## 3. Results

### 3.1. Aspein is predicted as a fully disordered protein prone to phase separation

From the N- to the C-terminus, the Aspein protein is divided into four main domains **(Figure S2A)**: i) a cleaved signal peptide (res. 1-19), ii) a more hydrophobic N-terminal (Nter) domain (res. 20-46), followed by iii) a short polyDA or DA stretch (res. 47-77), and terminated by iv) an extensively long polyD region consisting of repetitive SG(D)n blocks of variable Dn lengths, ranging from two to ten, and referred to as the SG(D)n domain (res. 78-413), accordingly. The mature protein is therefore 394 residues long for an averaged molecular mass of 39.31 kDa **(Figure 1A)**. Composition-wise, such sequence organisation results in a dramatic and unusual enrichment in Asp (60.4%), Ser (13.2%), and Gly (16.0%) amino acids **(Figures 1A and S2B)**. This exceptionally acidic nature is further exacerbated by the total absence of both positively charged arginine (Arg, R) and lysine (Lys, K), as well as pH-sensitive histidine (His, H) sidechains.

**Figure 1.**
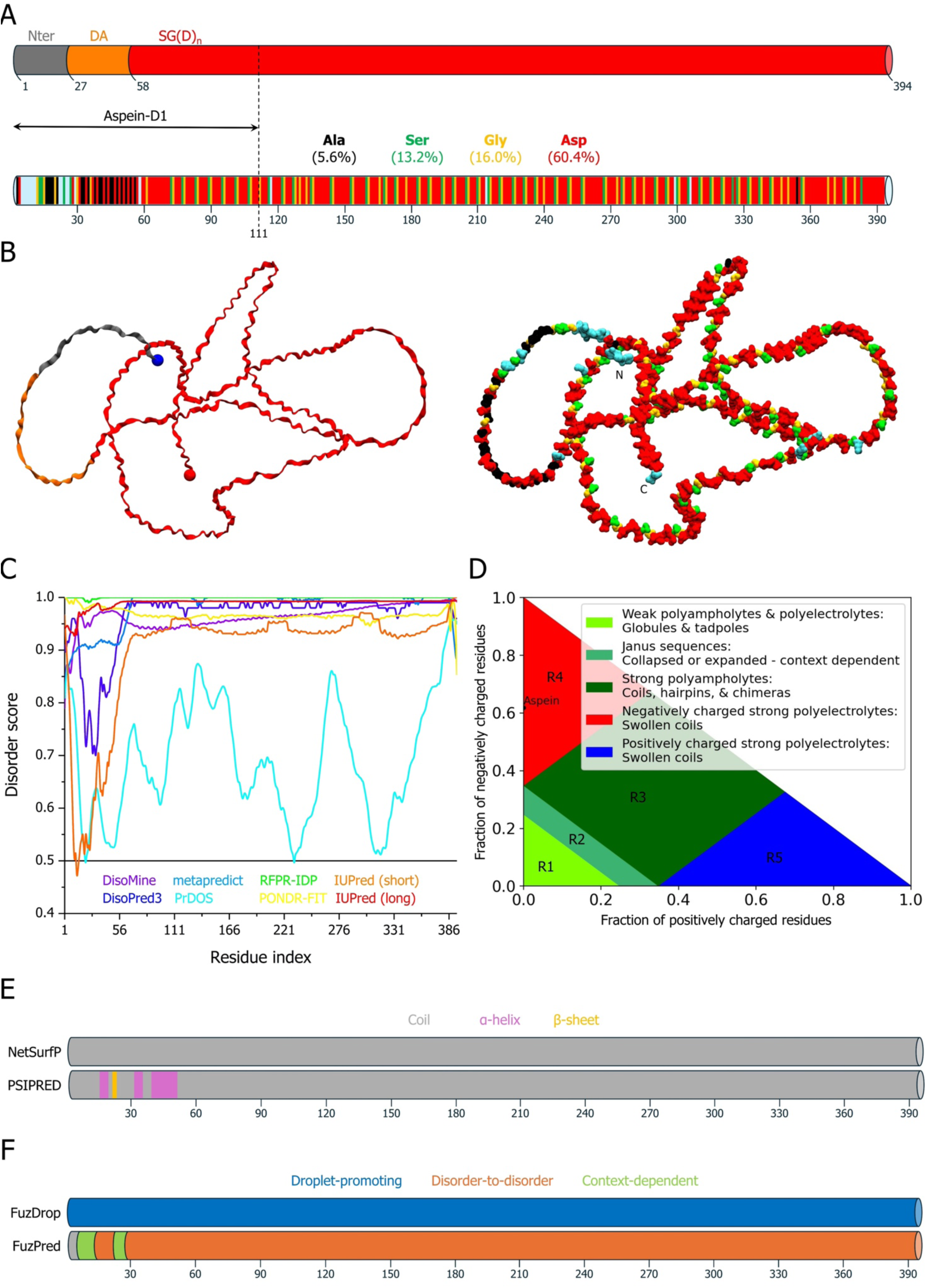
Sequence-based features and intrinsic disorder and phase separation predictions of Aspein. (A) Sequence domain organisation (Nter, DA, SG(D)_n_) and amino acid composition (Ala, Ser, Gly, Asp) of the mature full-length protein. The experimentally validated truncated version (Aspein-D1) is illustratively indicated. (B) Structural MMseqs2-AlphaFold2 model represented in ribbon (left) and van der Waals surface (right) representation and coloured at the domain and amino acid level according to the code used in panel A. N- and C-termini are either highlighted as blue and red spheres (left), respectively, or explicitly written (right). (C) Disorder propensity plots generated along the full-length protein sequence with eight independent predictors (DisoMine, DisoPred3, metapredict, PrDOS, RFPR-IDP, PONDR-FIT, IUPred-short, IUPred-long). (D) Das-Pappu state diagram of IDP conformational ensembles. (E) Secondary structure (NetSurfP, PSIPRED), as well as (F) LLPS (FuzDrop) and mode of binding (FuzPred) propensities projected on the schematised full-length protein sequence.

Another striking feature is the lack of aromatic residues, at the exception of one phenylalanine (Phe) at the N-terminus. In that quality, Aspein is not expected to absorb visible or near-UV light or show noticeable emission in the UV range, thus limiting its detection and characterisation *in vitro*. Furthermore, Aspein is expectedly unable to oligomerise through disulphide bridging as it is devoid of cysteine (Cys).

From a structural point of view, the resulting AF2 model adopts an expanded random coil conformation regardless of the domain **(Figure 1B)**, suggesting a dominant unfolded state, i.e. IDP. This character reasonably arises from electrostatic repulsions within the chain **(Figure S3B)**. Eukaryotic IDPs are statistically biased towards higher Glu content, assumed to induce helicity in disorder-to-order transitions occurring upon interaction.^85^ Therefore, the strong aspartic character of Aspein presumably helps maintaining a highly disordered state in the bound state, which could be relevant to its function. In that sense, sequence-based predictions and averaged intrinsic disorder (PPID∼99%) unambiguously designate Aspein as a fully disordered protein **(Figure 1C)**, with only a slight ordering proneness in the Nter domain arising from a higher hydrophobic character **(Figure S3A)**. Resulting from the combination of low hydropathic and highly charged characters, both the CDF **(Figure S3C)** and CH **(Figure S3D)** plots corroborate the IDP classification. Complementarily, all the AF2-generated models coherently display very low predicted confidence scores all along the sequence, which also advocates for a highly unfolded state **(Figure S3E)**.

Based on the fractions of positively and negatively charged residues, the Das-Pappu phase diagram discriminates IDP conformational ensembles according to five main groups.

In good agreement with the protein charge distribution **(Figure S3B)**, CIDER unsurprisingly assigns Aspein to the R4 region, comprising strong polyanionic sequences existing as extended conformers or swollen coils, similar to a pre-molten globule state **(Figure 1D)**. The very low values of κ (0.16) and Ω (0.12) charge patterning parameters also reflect such structural preference. Accordingly, secondary structure predictions do not reveal any significant propensity for structuring elements since all the sequence is primarily identified as a continuous statistical coil **(Figure 1E)**. Such property is also captured by the protein binding mode, engaging most exclusively into disorder-to-disorder or fuzzy interfaces **(Figure 1F)**, as well as by the absence of MoRFs detected in the sequence. Indeed, MoRFs are short and disordered segments that undergo disorder-to-order transitions upon interaction. Finally, Aspein is expected to be susceptible to LLPS driven by its two acid-rich polyDA and polyD LCRs defined by the DA and SG(D)n domains, respectively **(Figure 1F)**. Three out of four LLPS-oriented predictors, i.e. FuzDrop (pLLPS = 1.00), deePhase (pLLPS = 0.78), and MolPhase (pLLPS = 0.99), converge towards a high condensation propensity.

Considering its intrinsic disorder and functional role in the formation of mineralised organic frameworks, we propose that Aspein shall be considered as an **intrinsically disordered organic matrix protein (IDOMP)** from this point forward. As such, the coined IDOMP group should not only be used to encompass disordered SMPs but also other IDPs and proteins with IDRs that actively participate in biomineralisation processes.

### 3.2. Aspein is a fuzzy and selective ionic binder promoting calcite formation

Aspein is able to precipitate calcite *in vitro* and its Aspein-D1 version (res. 1-111), comprising the Nter and DA domains, as well as the first repeats of the SG(D)n region **(Figure 1A)**, was shown to reproduce the behaviour of the full-length protein.^35^ However, the precise mechanisms driving the CaCO_3_ polymorph selection remains uncharted. Therefore, we aim at deciphering the interplay of its conformational ensemble, amino acid composition, and biominerals formation by simulating a structural model of Aspein-D1 in different ionic environments while considering its intrinsic disorder and the aragonitic conditions of modern seawater. Protein structure and ion properties in 10 mM NaCl, as a monovalent control, were compared with the 10 mM CaCO_3_ and 10 mM CaCO_3_ + 50 mM MgCl_2_ divalent systems, the latter corresponding to the concentrations encountered in *P*. *fucata* extrapallial fluid where Aspein is secreted.

RMSD distributions show that the protein conformational ensemble is more heterogeneous in NaCl than for the two CaCO_3_ conditions **(Figure 2A)**. Populations distributed at lower RSMD values indicate that increased homogeneity is further observed without MgCl_2_. Those discrepancies are exemplified by clustering and PCA analysis **(Figure S4)**. Whereas in NaCl the protein occupies a diffuse and uncorrelated conformational space, CaCO_3_ induces Aspein to converge to more well-defined conformations, as shown by more correlated motions and the presence of defined basins on the PC projection. Instead, in agreement with the RMSD profile, the presence of MgCl_2_ nonetheless delays the cluster convergence. Such structural discrepancies are accompanied by a significant gain in compactness **(Figure 2B)** and desolvation **(Figure 2C)** of the chain upon CaCO_3_ addition, regardless of the presence of Mg^2+^ cations. The latter nonetheless lead the protein to adopt slightly more expanded conformations. As such, cation divalency appears to specifically trigger the collapse of the Aspein structure. The strong difference between NaCl and CaCO_3_ conditions expectedly induces a distinct reorganisation of water solvation shell around the protein **(Figure 2D)**. Indeed, via chain extension in NaCl, Aspein remains highly hydrated, thus increasing the water density particularly close to its highly solvent-exposed surface. On the contrary, divalent cations drive a chain collapse that expels water molecules to farther hydration shells, explaining the density increase observed at longer range. Once more, only subtle changes distinguish the two CaCO_3_ simulations, as further density shift and less variability between replicates characterise the Mg-free system. Furthermore, water dynamics and (de)ordering at the protein surface are also indicators of ion binding events.^86^

**Figure 2.**
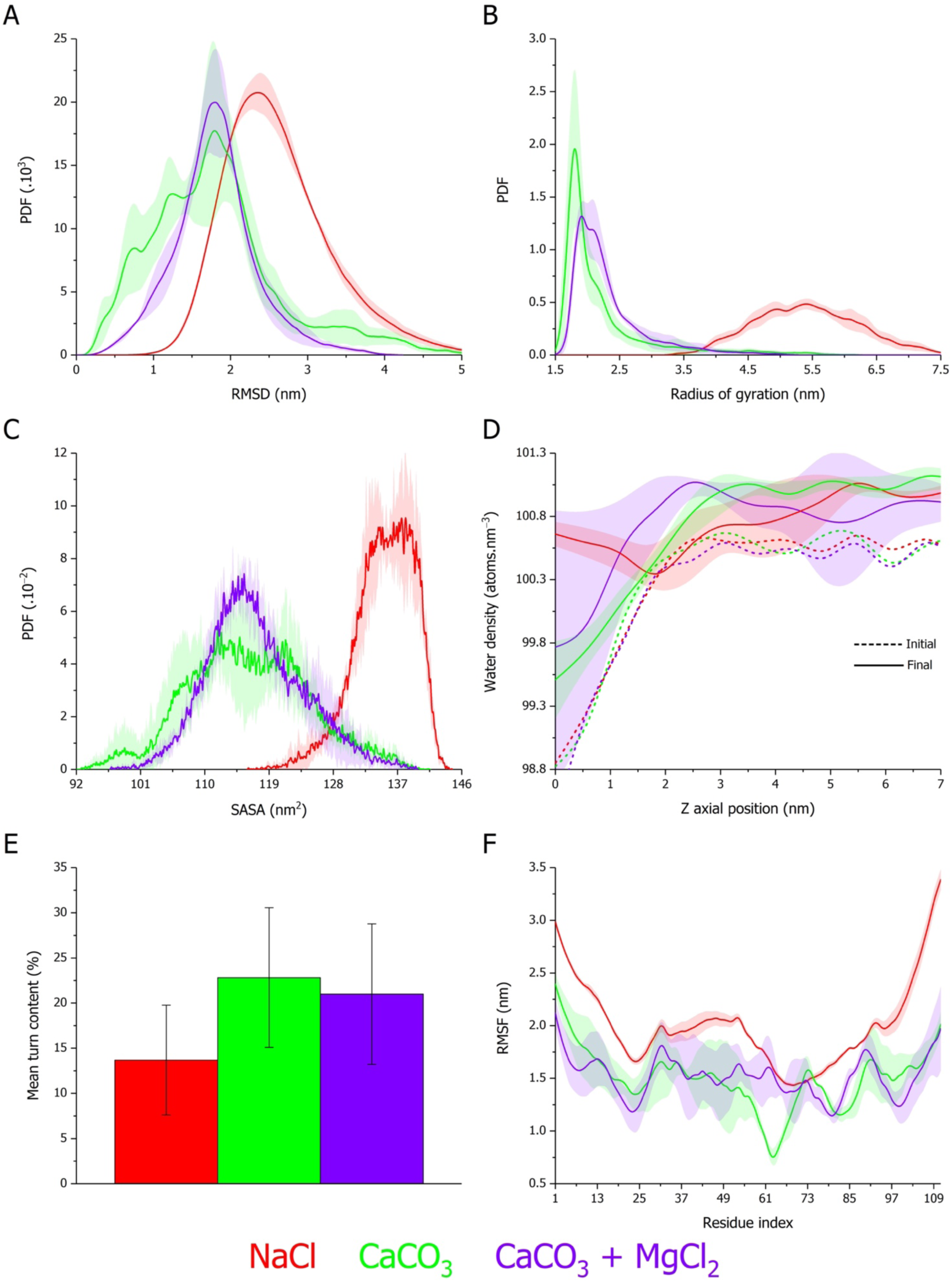
Structural and hydration properties of Aspein in different ionic environments. Probability density function (PDF) of (A) root-mean square deviation (RMSD), (B) radius of gyration (R_g_), and (C) solvent accessible surface area (SASA) of Aspein-D1 simulated for 1 µs in 10 mM NaCl (red), 10 mM CaCO_3_ (green), and 10 mM CaCO_3_ with 50 mM MgCl_2_ (purple). (D) Water atoms density relative to the protein geometric center and projected along the Z axis box dimension at the initial (dashed line) and final (solid line) stages of the simulation. (E) Time-averaged turn content and (F) backbone root-mean square fluctuation (RMSF). Curves or histograms correspond to the average of triplicates with the standard deviation represented as a trace (shaded area) in the condition-associated colour or error bars, respectively.

Intriguingly, despite such dramatic conformational transition, the protein predominantly remains in a disordered statistical coil state amongst all the conditions **(Figure S5)**. However, while this is particularly true in NaCl **(Figures S5A-B)**, a clear increase in turn content, which arises and to some extent oscillates over time, is comparatively observed for the other two systems **(Figures 2E and S5C-F)**. Interestingly, turn enrichment mainly occurs in the SG(D)n domain, in which the aspartic density is at its highest. Although secondary structural elements of α and β character are detected, their sporadic generation is not statistically significant and confined to the more hydrophobic N-terminal domain **(Figure S5)**. In addition, since such residual motifs were also formed in the Nter region in NaCl, their dependence on the presence of Ca^2+^ or Mg^2+^ ions can be ruled out. The permanence of unrestricted and flexible coil conformations is reflected by the RMSF profile of Aspein in the presence of Na^+^ ions **(Figure 2F)**. RMSF values are overall lower with CaCO_3_ showing that ionic divalency promotes the protein chain collapse and restriction of the conformational mobility seemingly through ion binding. However, each domain locally exhibits dissimilar behaviours: i) steady flexibility decrease along the Nter domain; ii) persistence of a high flexibility for the DA domain; iii) backbone stiffening for the SG(D)n polyD IDR with further local reductions. The latter are coherent with the increased turn content in the SG(D)n region, applying structural constraints, and likely driven by the interactions involving Asp residues with Ca^2+^ and/or Mg^2+^ cations. The addition of MgCl_2_ on the contrary enhances the flexibility of the segment spanning the DA and SG(D)_n_ domains (res. 50-71), suggesting a locally looser conformer albeit a higher ionic concentration. This is supported by the per-domain ΔR_g_ and ΔSASA evolution over the course of the simulations **(Figure S6)**. Indeed, the DA domain is mostly affected by the introduction of MgCl_2_, slightly diminishing its compaction and achieving a partially dehydrated state close to that of the initial structure. Otherwise, all trends are similar between the CaCO_3_ conditions with and without MgCl_2_, showing that the protein as a whole rapidly and considerably decreases its Rg and SASA chiefly through the collapse and desolvation of the SG(D)_n_ domain. This compact conformation persists from 200 ns onwards, which is an important difference compared to the other IDRs that remain more dynamic. Consistent with the data reported so far, the protein chain rather expands in NaCl, and only the Nter becomes partly compacted and dehydrated, presumably due to its enhanced hydrophobic character.

By examining the properties of the different ionic species present in the simulated systems, the structural changes of Aspein described so far are remarkably driven by selective protein-ion interactions. Mean square displacement (MSD) is a useful parameter for tracking and appreciating the diffusion characteristics of individual ions in a medium **(Figures 3A-B)**.^87^ The linear MSD increase experienced by Cl^−^ anions indicates that they obey a Brownian motion, and hence are not interacting with Aspein. Instead, the motion of Na^+^ and Mg^2+^ cations are to some extent impeded, as shown by the corresponding lagging MSD curve. Weak, nonspecific, and dynamic binding of these cations to Aspein may be responsible for their anomalous subdiffusion behaviour. On the other hand, both Ca^2+^ and CO_3_^2−^ ions concomitantly converge to a stable average whatever the presence or absence of Mg^2+^. Since plateauing MSD profiles indicate the confinement of particles, the restriction of ionic motion for Ca^2+^ and CO_3_^2−^ imply that they engage into static and specific interactions with Aspein. The calculation of the number of bound ions within a 4 Å cut-off around the protein surface confirms this fact **(Figures 3C-D)**. Indeed, compared to other ions, a small fraction of which is sporadically found in the vicinity of the protein, Aspein preferentially binds to Ca^2+^ and CO_3_^2−^ with a gradual enrichment up to saturation. This tendency is even more striking considering the high Mg/Ca ratio used in the CaCO_3_ + MgCl_2_ simulations. It is also unambiguously ascertaining the selective nature of Aspein. Time-dependently, Ca^2+^ are the first ions to crowd around the protein before attracting CO_3_^2−^ anions to the Ca^2+^-bound surface, in a sort of electrostatic double layer. Whilst such stepwise addition is unaltered by the presence of Mg^2+^ cations, the latter reduce the number of those ions in the bound state **(Figure 3D)**.

**Figure 3.**
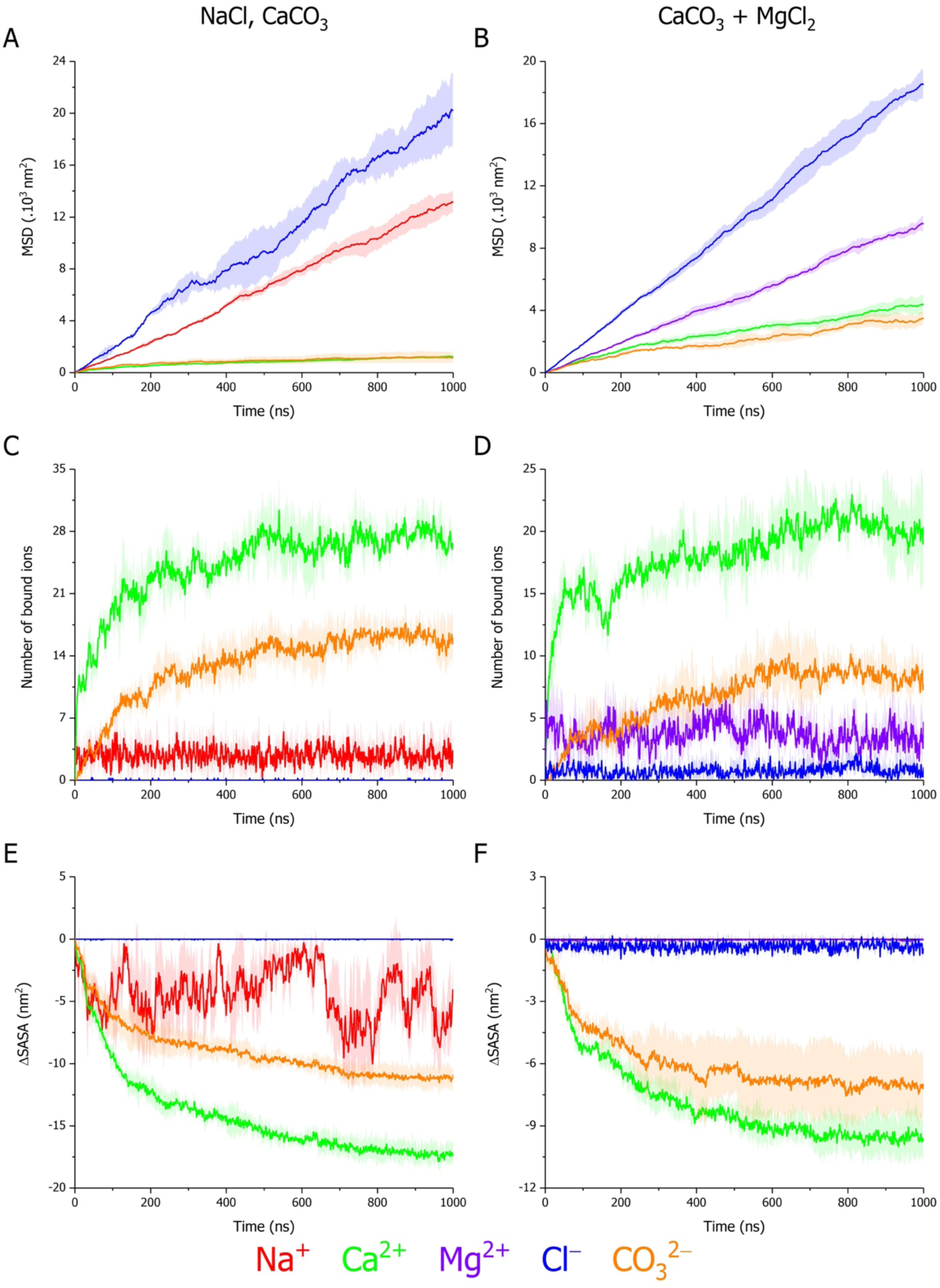
Aspein selective effects on the diffusion, binding, and solvation of ionic species. Time evolution of (A-B) mean square displacement (MSD), (C-D) number of contacts within a 4 Å cut-off around the protein surface, and (E-F) solvent accessible surface area variation (ΔSASA) relative to the first timestep of Na^+^ (red), Ca2^+^ (green), Mg^2+^ (purple), Cl^−^ (blue), and CO_3_^2−^ (orange) ions encountered in the Aspein-D1 systems simulated for 1 µs in (A, C, E) 10 mM NaCl (left), 10 mM CaCO_3_ (left), and (B, D, F) 10 mM CaCO_3_ with 50 mM MgCl_2_ (right). On each panel, curves correspond to the average of triplicates with the standard deviation represented as a trace (shaded area) in the ion-associated colour.

From a domain-oriented point of view, both polyD domains, i.e. DA and SG(D)_n_, unsurprisingly participate in Ca^2+^ binding with a higher Ca-bound content for SG(D)_n_ due to the difference in Asp density **(Figure S7)**. Furthermore, Ca^2+^ and CO_3_^2−^ are the only tested ionic species to experience a progressive and converging desolvation over time **(Figures 3E-F)**. In light with the dehydrated chain collapse and motion confinement that Aspein and Ca^2+^/CO_3_^2−^ ions undergo, concomitant desolvation of the latter can be explained by the ions being trapped into the collapsed structure of the protein, especially in the SG(D)_n_ domain, and stabilised into CaCO_3_ aggregates.

In that respect, the existence of protein-bound CaCO_3_ phases is first verified in our simulations by comparing the proximal radial distribution function (pRDF) of ions with respect to Asp residues. As expected from the existence of direct electrostatic contacts between Asp carboxylates and Ca^2+^ cations, a strong peak is detected at ∼2.3 Å **(Figures S8A-B)**. Although the first peak associated with Mg^2+^ is located at ∼1.8 Å, such a nearer position originates from its smaller ionic radius compared to Ca^2+^. In addition, its intensity is relatively weaker in regard to the used Mg^2+^ concentration and the height of the Ca^2+^ peak. Interestingly, CO_3_^2−^ anions are not randomly partitioned, as their main peak, appearing at ∼3.8 Å, is indicative of their positioning to the second interaction shell of Asp amino acids. The second, but weaker, Ca^2+^ peak at ∼4.2 Å further advocates for the stabilisation and compartmentalisation of CaCO_3_ by Asp residues. By arbitrarily isolating D98 and D93 in the CaCO_3_ boxes without and with Mg^2+^, respectively, we see indeed that such organisation extends at longer range with the alternation of well-defined Ca^2+^ and CO_3_^2−^ peaks **(Figure S8C)**. Moreover, CaCO_3_ ordering occurs with time, as Ca^2+^ and CO_3_^2−^ ions are disorganised when they bind the protein at the early phases of the simulations. While Mg^2+^ cations do not confine into ordered structures as a result of their interaction with Aspein **(Figure S8E)**,^88,89^ they partially disrupt the CaCO_3_ organisation, especially at long range **(Figure S8D)**, which could originate from the reduction of the Asp fraction available for Ca^2+^ binding.

Secondly, preferential Ca-CO_3_ arrangements have also been analysed to decipher whether they echo the selective polymorph precipitation reported *in vitro* for Aspein **(Figure 4)**. The distance distribution between Ca^2+^ ions and C atoms in carbonate allows to determine the coordination shell of Ca^2+^, i.e. mono- (second peak) or bidentate (first peak) carbonate, depending on the contribution of one or two oxygen atoms, respectively.^90^ Whereas vaterite and aragonite comprise a mix of mono- and bidentate ligands, calcite is exclusively monodentate, which is expected to be the most stable.^91–93^ As for aragonite, ACC presumably contains a statistical 1:1 monodentate:bidentate ratio, although several studies pinpointed a significant monodentate enrichment (∼70-80%) in the amorphous phase.^94,95^ Remarkably, Aspein induces an overtime monodentate enrichment, Mg^2+^ cations having no influence on this arrangement **(Figures 4A-B)**. Such fact suggests not only the stabilisation of ACC but also the time-dependent formation of calcite clusters. The emergence of calcitic features from ACC is also substantiated by the pRDF of Ca^2+^ with respect to the oxygen atoms of CO_3_^2−^ **(Figures 4C-D)**.^96–99^ The first peak at ∼2.2 Å corresponds to ACC or calcite, whereas it is typically shifted to ∼2.4-2.5 Å for aragonite and vaterite polymorphs.^98,100^ Once again, the presence of Mg^2+^ has little to no impact on calcite generation and, more importantly, does not trigger the appearance of peaks associated with aragonite **(Figure 4D)**. Interestingly, a peak corresponding to vaterite is detected in both conditions and could be relevant with respect to the crystallisation pathway of CaCO_3_, as vaterite-like intermediates are known to be induced by long polyD polymers through pseudomorphic transformation before a calcitic switch via particle attachment.^101,102^ The pRDF profiles of Ca-Ca correlation show a similar preference for ACC and calcite regardless of the Mg^2+^ excess **(Figures 4E-F)**.^96–98,100,103^ Nonetheless several vaterite features are still detected, but those of calcite become prevalent with time, thus reflecting the gradual ordering of protein-captured CaCO_3_ clusters into crystalline calcite nuclei. Once more, the coexistence of vaterite remains mechanistically relevant for the precipitation pathway of calcite.^101,102^

**Figure 4.**
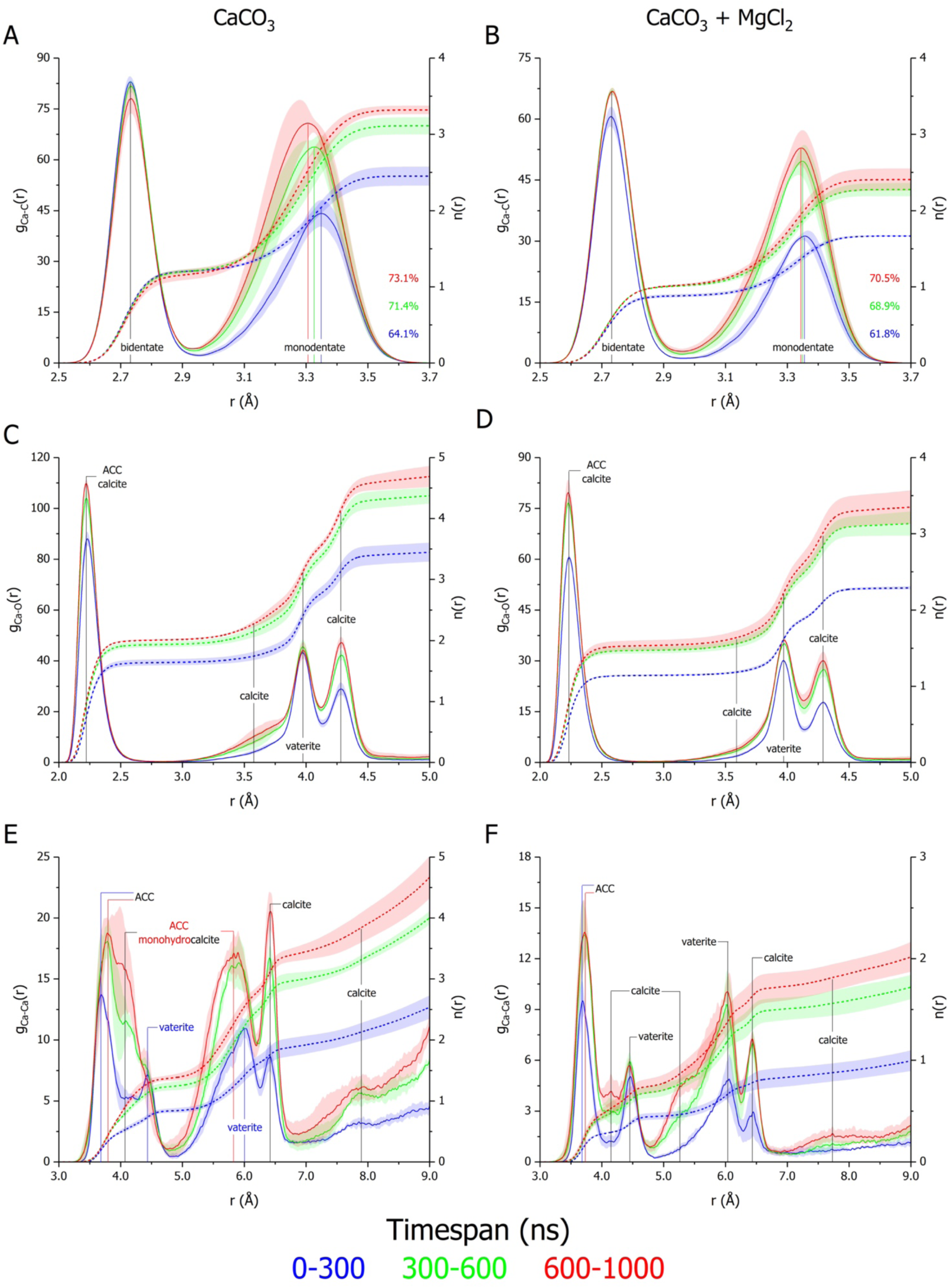
Aspein selective effects on CaCO_3_ polymorphism in extrapallial conditions. Proximal radial distribution function (pRDF) g(r) (solid line) and cumulative number count n(r) (dashed line) of (A-B) Ca-C, (C-D) Ca-O, and (E-F) Ca-Ca atoms from Ca^2+^ and CO_3_^2−^ ions encountered in the Aspein-D1 systems simulated for 1 µs in (A, C, E) 10 mM CaCO_3_ (left) and (B, D, F) 10 mM CaCO_3_ with 50 mM MgCl_2_ (right). Trajectories are divided into three subsequent timespans, i.e. from 0 to 300 (blue), 300 to 600 (green), and 600 to 1000 ns (red), to capture the evolution of g(r) and n(r) over time. On the g_Ca-C_(r) panels (A-B), percentages are indicative of monodentate population determined from n(r) ratios. On the g_Ca-O_(r) and g_Ca-Ca_(r) panels (C-F), detected pRDF peaks are labelled according the CaCO_3_ polymorph they are associated with. On each panel, curves correspond to the average of triplicates with the standard deviation represented as a trace (shaded area) in the timespan-associated colour.

Through the combination of the *in silico* data presented so far with snapshots at chosen timeframes, we illustrate how Aspein specifically captures CaCO_3_ and selects calcite, even in Mg-enriched media **(Figure 5)**. Indeed, the protein responds differently towards cationic environments **(Figure 5A)**. Whereas Aspein preferentially occupies a highly flexible, extended, and hydrated state due to the insufficiency of Na^+^ cations to screen Asp-Asp intrachain repulsion, Ca^2+^ tightly and rapidly binds to the polyAsp domains driving an overall chain desolvation and structure-less collapse trapping and stabilising ions via the formation of Ca^2+^-bridged turns **(Figure 5B)**. Attracted by the charge density harboured by the Ca-bound Aspein, CO_3_^2–^ anions incorporate into the structure in which Asp residues are responsible for stabilising ACC and calcite phases. Indeed, distinctive organisational features of calcite are observed by the end of the simulations regardless of elevated Mg^2+^ levels in the direct environment of Aspein **(Figure 5B)**.^100,104^ While adopting highly extended structures due to electrostatic repulsion in its free state, the conformational ensemble of Aspein can be more accurately described as fuzzy in the presence of CaCO_3_, as the chain retains its dynamical properties without significant disorder-to-order transitions upon binding to its ionic targets.^105^ Furthermore, Aspein compartmentalises ionic species via its Asp sidechains that not only catch Ca^2+^ ions and sequester CaCO_3_ clusters within the protein collapsed structure but also prevent Mg^2+^ cations to integrate ACC by cloistering them at the protein surface. All these effects noteworthily contribute to the selection of calcite over aragonite in the extrapallial fluid of *P. fucata*.

**Figure 5.**
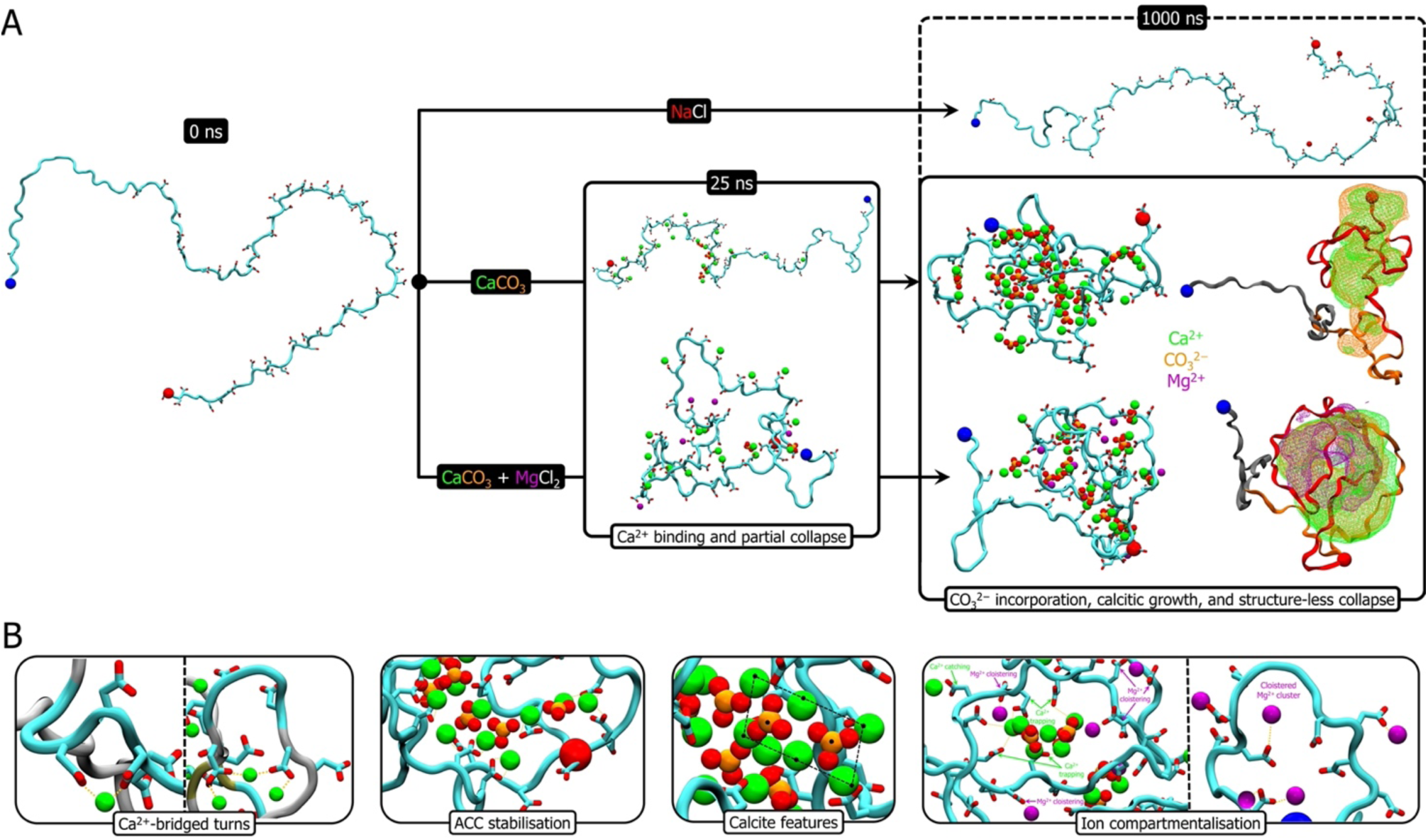
– Structural and molecular insights into Aspein-mediated CaCO_3_ crystallisation. From representative trajectories, selected snapshots illustrate (A) the time-dependent conformational response of Aspein to different ionic environments, as well as (B) the diversity of roles exerted by Asp amino acids for CaCO_3_ sequestering and polymorph selection. On the different panels, Na^+^ (red), Ca^2+^ (green), Mg^2+^ (purple), CO_3_^2−^ (orange and red for C and O atoms, respectively) ions found within a 4 Å cut-off around the protein surface are depicted as spheres using scaled up van der Waals radii, while the Aspein chain (cyan) is displayed in cartoon representation with the N- and C-termini respectively pinpointed as blue and red spheres, Asp residues in licorice representation (red for Oδ atoms) and cation-Asp interactions as yellow dashes. Time-averaged volumetric density maps (1000 ns panel, far right) of Ca^2+^ (green), Mg^2+^ (purple), and CO_3_^2−^ (orange) ions are represented as a wireframe surface in the ion-associated colour, as well as superimposed on the final Aspein structure coloured according to its Nter (dark grey), DA (dark orange), and SG(D)_n_ (dark red) domains showed in ribbon.

### 3.3. Aspein condensates into Mg-free protein-CaCO_3_ clusters via its SG(D)_n_ motifs

Biogenic crystallisation has recently been described in the context of LLPS and different predictions agree on the high phase separation propensity of Aspein. To better understand its environment-dependent condensation behaviour, we restricted the system to its two most abundant ion binding motifs, i.e. SG(D)3 and SG(D)4, and mixed twenty all-atom copies of each in the previously tested saline conditions.

With respect to the control NaCl condition, CaCO_3_ induces a more homogeneous, compacted and dehydrated state of the peptide chains **(Figures 6A-D)**. Such compaction seemingly arises from their clustering with ions, as suggested by their confined MSD profile and restricted backbone flexibility **(Figures 6E-F)**. The introduction of a high concentration of Mg^2+^ cations leads to an intermediate state which likely reflects an altered condensed form or a modified self-assembly pathway. Such claims are supported by the behaviour of the different ionic species, the tendencies of which are echoing those described for the Aspein-D1 single chain system. Whilst Mg^2+^ cations follow a subdiffusion pattern and their concentration does not increase in the vicinity of the chains, specific interactions engage the SG(D)3-4 peptides with Ca^2+^ and CO_3_^2–^ ions that are progressively desolvated and confined upon binding, even upon the addition of excess MgCl_2_ **(Figures 7A-F)**. As for Aspein-D1, the very first binding event involves Ca^2+^ cations followed by the attraction of CO_3_^2–^ anions, meaning that SG(D)3-4 peptides do not interact with preformed CaCO_3_ clusters **(Figures 7C-F)**. Moreover, the resulting proteinaceous assemblies, while shielding from Mg^2+^ cations and sequestrating CaCO_3_, remain in a fuzzy state, as no secondary structure generation can be observed in the three tested conditions **(Figure S9)**.

**Figure 6.**
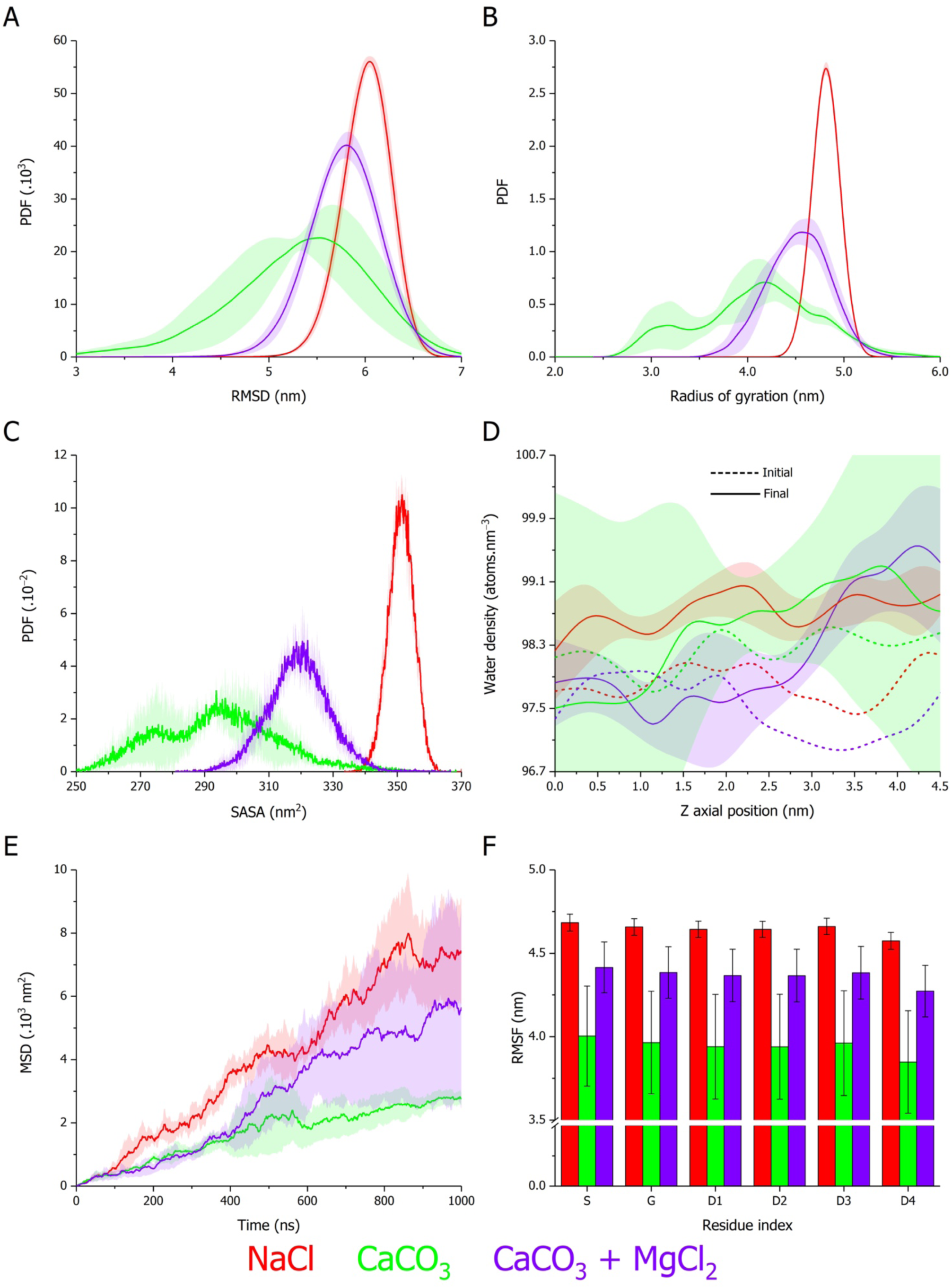
Structural and hydration properties of a SG(D)_3-4_ peptides mixture in different ionic environments. Probability density function (PDF) of (A) root-mean square deviation (RMSD), (B) radius of gyration (R_g_), and (C) solvent accessible surface area (SASA) of mixed SG(D)_3_ and SG(D)_4_ free monomers (twenty copies each) simulated for 1 µs in 50 mM NaCl (red), 50 mM CaCO_3_ (green), and 50 mM CaCO_3_ with 250 mM MgCl_2_ (purple). (D) Water atoms density relative to the peptide chains geometric center and projected along the Z axis box dimension at the initial (dashed line) and final (solid line) stages of the simulation. (E) Time evolution of peptide backbone mean square displacement (MSD). (F) Time-averaged backbone root-mean square fluctuation (RMSF) determined per residue across the forty dispersed chains. Curves or histograms correspond to the average of triplicates with the standard deviation represented as a trace (shaded area) in the condition-associated colour or error bars, respectively.

**Figure 7.**
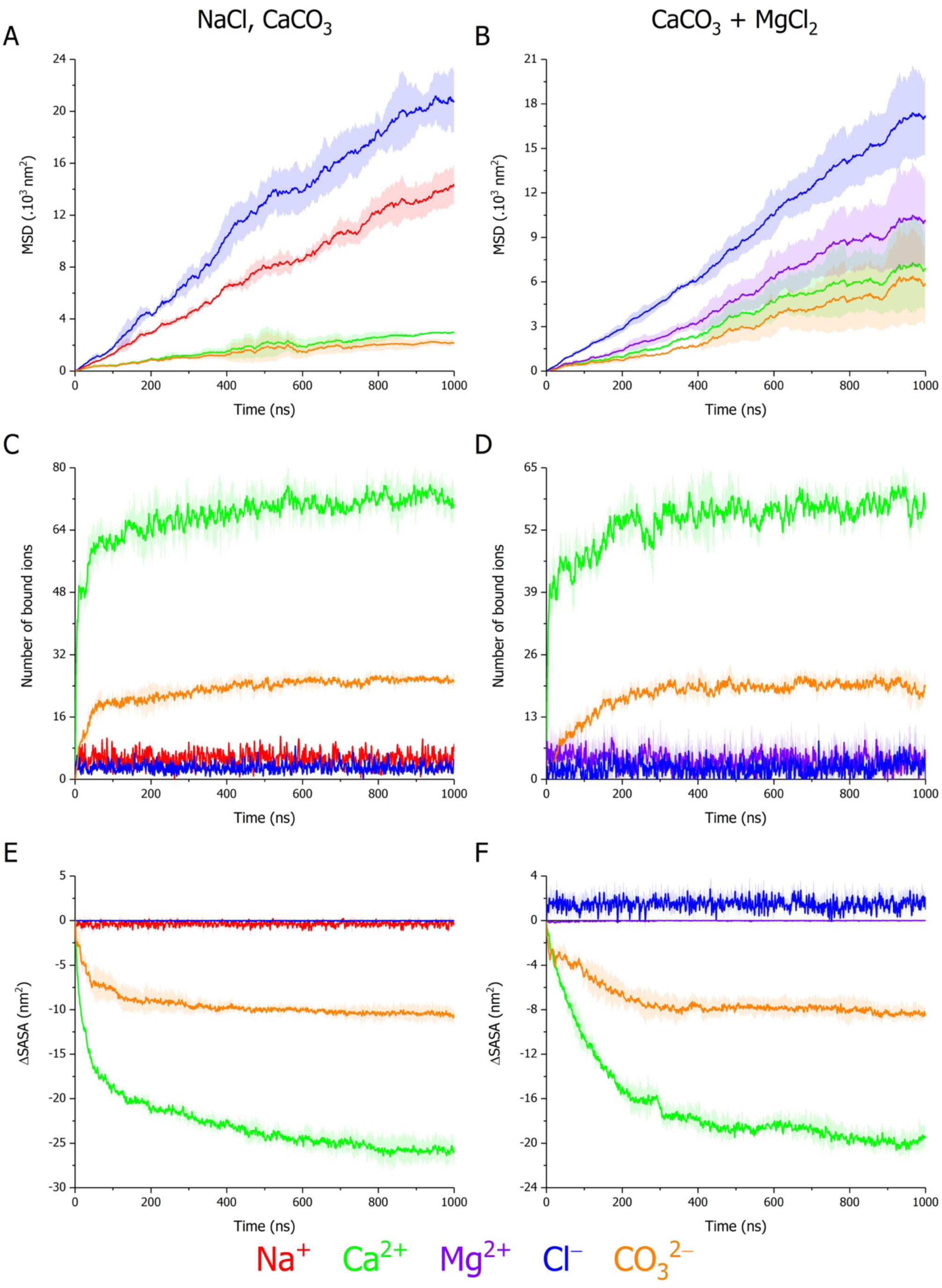
Selective effects of the SG(D)_3-4_ peptides mixture on ion diffusion, binding, and solvation. Time evolution of (A-B) mean square displacement (MSD), (C-D) number of contacts within a 4 Å cut-off around the peptides surface, and (E-F) solvent accessible surface area variation (ΔSASA) relative to the first timestep of Na^+^ (red), Ca2^+^ (green), Mg^2+^ (purple), Cl^−^ (blue), and CO_3_^2−^ (orange) ions encountered in the mixed SG(D)_3_ and SG(D)_4_ free monomers (twenty copies each) systems simulated for 1 µs in (A, C, E) 50 mM NaCl (left), 50 mM CaCO_3_ (left), and (B, D, F) 50 mM CaCO_3_ with 250 mM MgCl_2_ (right). On each panel, curves correspond to the average of triplicates with the standard deviation represented as a trace (shaded area) in the ion-associated colour.

To further attest the CaCO_3_-driven phase separation of Aspein, we have scanned the atom or charge density along one of the system box axes to obtain a good descriptor of such spatially coordinated phenomenon.^106^ The comparison of the charge distribution between SG(D)3-4 Asp residues and Na^+^/Cl^−^ ions at the beginning and end of the simulations reveals that their localisation is uncorrelated, showing the persistence of their random dispersion across the box **(Figures 8A-B)**. On the contrary, Asp residues become concentrated in a defined region of the box, which is accompanied by their co-localisation with Ca^2+^ and CO_3_^2–^ions **(Figures 8C-D)**. Whereas the density of Mg^2+^ cations remains unaltered and unbiased even in the presence of a high Mg/Ca ratio, the correlated separation of Asp amino acids with Ca^2+^/CO_3_^2–^ ions appears broadened along the X dimension **(Figures 8E-F)**. Such patterns substantiate the formation of protein-CaCO_3_ co-condensates, which are partly disrupted or fragmented by Mg^2+^ cations in extrapallial conditions **(Figure 9)**.

**Figure 8.**
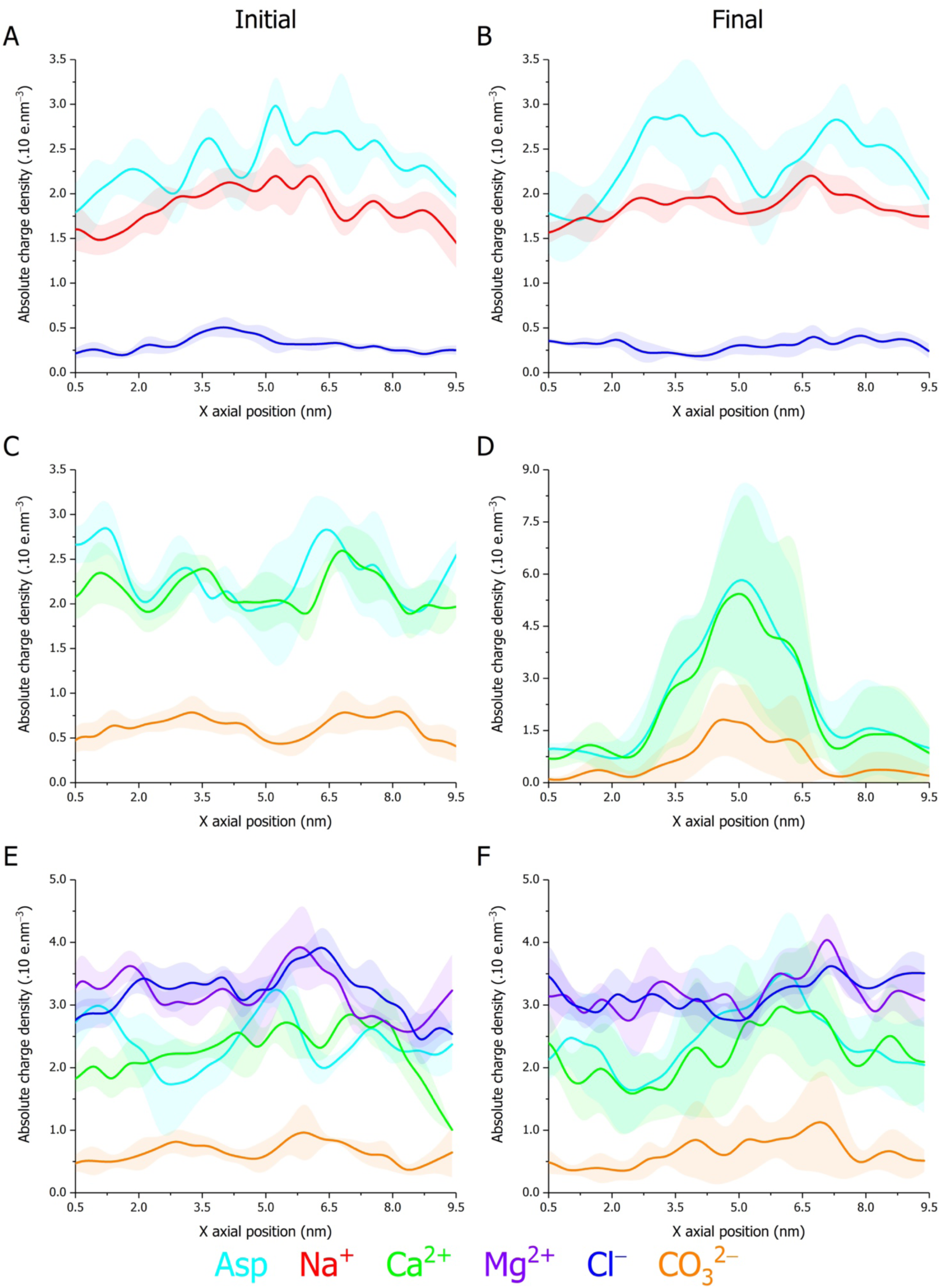
Selective and coupled SG(D)_3-4_ peptide-ion phase separation. Absolute charge density along the X axis box dimension at the initial (A, C, E) and final (B, D, F) simulation stages of Asp amino acids (cyan), as well as Na^+^ (red), Ca2^+^ (green), Mg^2+^ (purple), Cl^−^ (blue), and CO_3_^2−^ (orange) ions encountered in the mixed SG(D)_3_ and SG(D)_4_ free monomers (twenty copies each) systems simulated for 1 µs in (A-B) 50 mM NaCl, (C-D) 50 mM CaCO_3_, and (E-F) 50 mM CaCO_3_ with 250 mM MgCl_2_ (right). On each panel, curves correspond to the average of triplicates with the standard deviation represented as a trace (shaded area) in the ion-associated colour.

**Figure 9.**
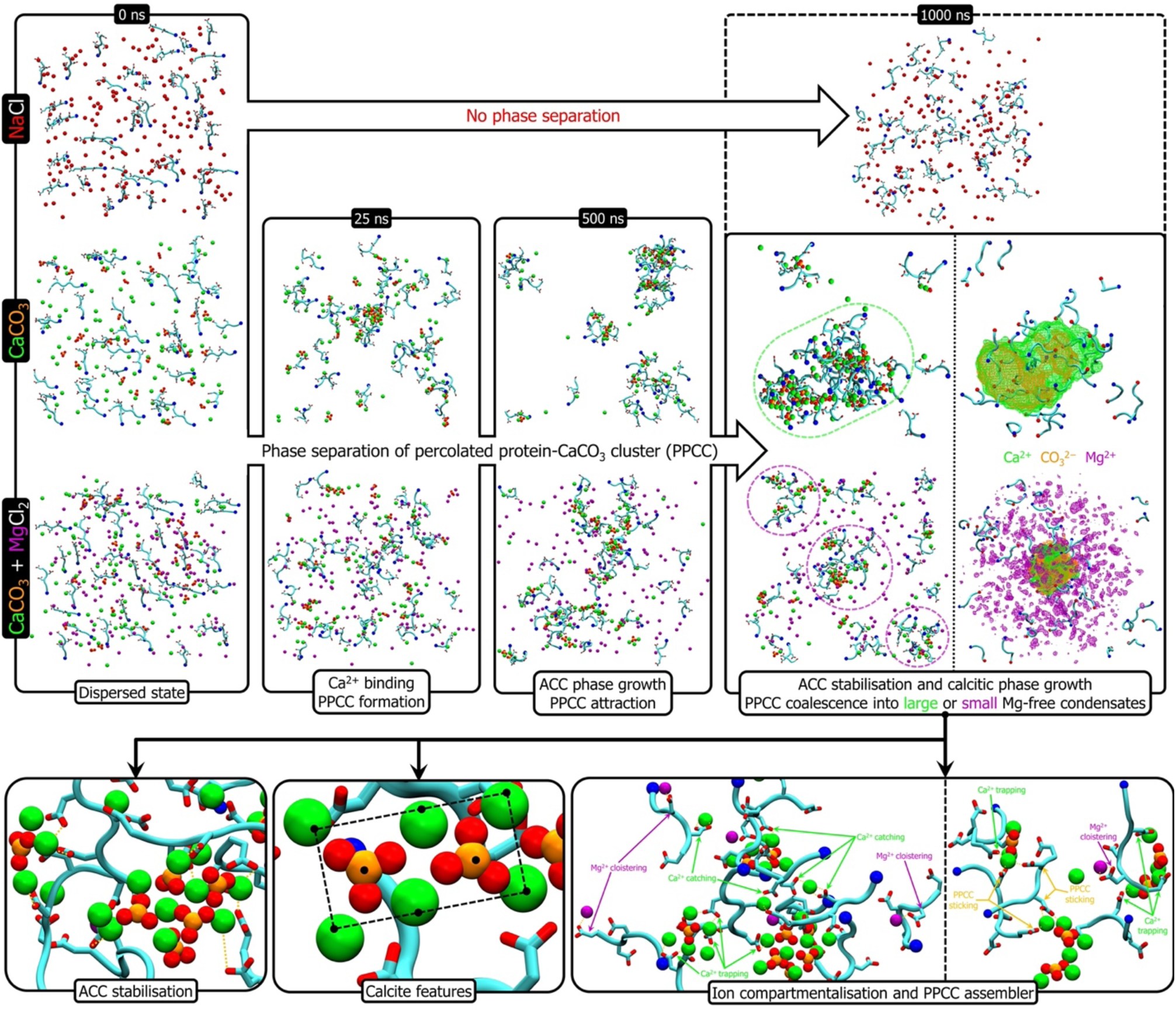
Molecular insights into Aspein-CaCO_3_ phase separation and its effects on CaCO_3_ crystallisation. From representative trajectories, selected snapshots illustrate the time-dependent phase separation behaviour of the SG(D)_3-4_ peptides mixture to different ionic environments (upper panel), as well as the diversified roles exerted by Asp amino acids for CaCO_3_ sequestering, polymorph selection, and PPCC condensation (lower panel). On the different panels, Na^+^ (red), Ca^2+^ (green), Mg^2+^ (purple), CO_3_^2−^ (orange and red for C and O atoms, respectively) ions are depicted as spheres using scaled up van der Waals radii, while the peptide chains (cyan) are displayed in cartoon representation with the N-terminus pinpointed as a blue sphere, Asp residues in licorice representation (red for Oδ atoms) and cation-Asp interactions as yellow dashes. Time-averaged volumetric density maps (1000 ns panel, far right) of Ca^2+^ (green), Mg^2+^ (purple), and CO_3_^2−^ (orange) ions are represented as a wireframe surface in the ion-associated colour, as well as superimposed on the final state of the SG(D)_3-4_ peptides mixture.

From dissecting the trajectories at several timesteps and visualising the corresponding snapshots, we may decipher and identify the environmental phase separation pathway of Aspein **(Figure 9)**. In a saline medium containing only NaCl, Aspein SG(D)3-4 domains are unable to phase separate and rather solvate as hydrated and extended monomers due to electronegative charge repulsion and their non-affinity for Na^+^ cations. Conversely, through their highly selective binding to Ca^2+^ cations, which screen the negative charges of Asp, Aspein chains undergo co-condensation in the presence of CaCO_3_, hence forming percolated protein-calcium carbonate clusters (PPCCs). Over the course of time, these PPCCs coalesce into larger protein-CaCO_3_ droplets thanks to free Asp carboxylate moieties acting as stickers that link Asp-stabilised CaCO_3_ phases.^107^

Although PPCC generation is not hindered upon the addition of excess Mg^2+^ cations, protein-CaCO_3_ condensates of smaller dimensions are formed. As such, Mg^2+^ cations partly shield inter-PPCC attraction. This fragmentation results from a fraction of peripheral Asp sidechains reallocated from PPCC coalescence into Mg^2+^ compartmentalisation at the droplets surface; otherwise, most Mg^2+^ cations remain solvated. Such response still allows for ACC and calcite stabilisation, while remarkably avoiding aragonite formation through Mg-ACC contamination. Indeed, features typical of ACC and calcite, even in the presence of MgCl_2_, are revealed by the examination of the pRDF profiles **(Figure S10)** that are also in good agreement with those of Aspein-D1 **(Figures 4 and S8)**: i) organised partition of Ca^2+^/CO_3_^2–^ ions **(Figures S10A-B)**, ii) enrichment in monodentate coordination **(Figure S10C)**, iii) overtime emergence of calcitic Ca-O and Ca-Ca peaks **(Figures S10D-E)**, and iv) absence of Mg embedding in CaCO_3_ phases **(Figure S10F)**.

Consistent with our previous observations, the unique aspartic density of Aspein enables its constituting Asp sidechains to endorse diversified roles that collaboratively participate in the selection of calcite in a pro-aragonite environment. In this framework, Asp residues either act as i) selective Ca^2+^ catchers to locally increase the concentration of Ca and attract CO_3_^2–^ anions, ii) Ca^2+^ trappers to stabilise CaCO_3_ phases or (pre)nuclei out of the bulk, iii) Mg^2+^ segregators to maintain these ions separated from CaCO_3_ clusters, thus preventing Mg poisoning that would favour aragonitic growth, or iv) stickers to assemble PPCCs into larger condensates.

## 4. Discussion

In light of our findings, we may validate the hypotheses formulated in 2008 by Takeuchi T., *et al*. regarding the Aspein-directed precipitation of calcite, while providing new insights into the interplay between IDOMPs and biogenic mineralisation.^35^ Takeuchi T., *et al*. speculated that Aspein would act as a selective Ca^2+^ accumulator in the high Mg^2+^ extrapallial conditions, the aggregation of ions being facilitated by the inherent physical flexibility of the protein SG(D)_n_ domain. Upon the encounter of Ca-bound Aspein with carbonate anions, the former would stabilise ACC before transforming into calcite. Indeed, our all-atom simulations have not only unveiled that Aspein specifically sequesters ACC transitioning into calcite nuclei over time but have also underscored the prominent roles of disordered regions and Asp amino acids in regulating the nucleation and polymorph selection processes **(Figure 10)**. As such, Aspein induces precipitation of calcite in the undersaturated CaCO_3_ conditions encountered in seawater by locally increasing the concentration of Ca^2+^ and subsequently trapping CO_3_^2−^ ions through protein structural collapse and phase separation, thus allowing to reach supersaturation. In addition, the nascent order of the calcite polymorph into which ACC will transform is already present in the CaCO_3_ clusters stabilised by the protein, both at the single molecule and multi-chain level.

**Figure 10.**
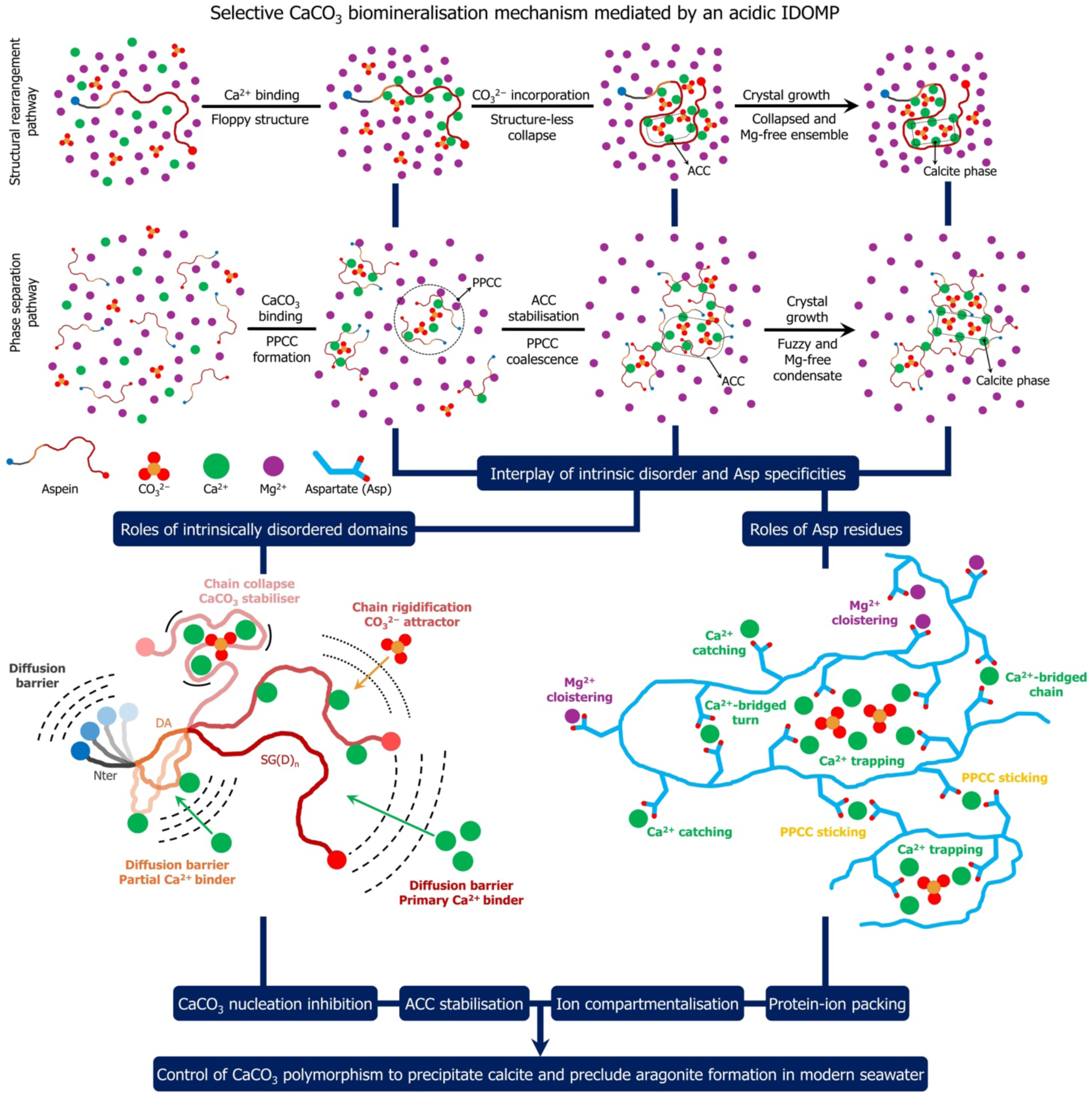
Aspein-mediated calcite biomineralisation mechanism. Schematised structural rearrangement and protein-ion phase separation pathways proposed to explain how Aspein, an unusually acidic IDOMP secreted in the aragonitic extrapallial fluid of *P*. *fucata*, selects calcite for the prismatic layer formation in light of the specific and concerted effects on CaCO_3_ diffusion, nucleation, saturation, and compartmentalisation exerted by its intrinsically disordered domains and unique Asp density. On the lower left panel, lines represent the evolution of the degree of chain flexibility from dynamic (dashes) to constrained (solid) for each Aspein domain upon ions incorporation. ACC, amorphous calcium carbonate; PPCC, percolated protein-CaCO_3_ cluster.

Polymorph selection is likely the result of several inhibitory effects on CaCO_3_ growth: i) inhibition of spontaneous nucleation of CaCO_3_ calcite from the bulk, which would result in elongated and sharp-edged crystals thus detrimental to the oyster; ii) inhibition of the dissolution-recrystallisation pathway by increasing the stability of Asp-bound ACC; iii) inhibition of the aggregation of CaCO_3_ clusters into vaterite nanospheres; iv) controlled generation of calcite nuclei.^101,102,108^ As such, calcite nucleation kinetics, while slowed down from bulk, likely remains faster than vaterite nucleation and aggregation through classical ion-to-ion attachment and/or secondary nucleation upon attachment and clustering of ACC particles. Notwithstanding the faster association observed between Aspein and Ca^2+^ cations, we cannot exclude that concomitant addition of CaCO_3_ pairs or small prenucleation clusters to PPCC occurs to some extent in our simulated model. Aspein could also stabilise calcite biocrystals by incorporating into the prismatic layer.

Furthermore, our MD simulations revealed that Aspein preferentially binds to, and stabilises pristine Ca-ACC, therefore preventing the adsorption of Mg^2+^ ions on ACC and the subsequent formation of aragonite. The protective role endorsed by this prismatic IDOMP is highly relevant with respect to a recent report disclosing that aragonite crystals actually embed Mg^2+^ cations in their structure through Ca/Mg substitutions and grow from Mg-trapping by ACC in seawater.^30^ This is also in very good agreement with the shell calcification mechanism experimentally suggested for another extremely acidic IDOMP, the Gh26 protein of the deep-sea mussel *Gigantidas haimaensis*, in Mg-enriched media.^27^ Complementarily, *P*. *fucata* has developed alternative strategies involving other IDOMPs to preclude aragonite crystallisation in the prismatic layer. One of such examples concerns the acidic SMP PfN44 that, contrary to Aspein, has a greater affinity for Mg^2+^ compared to Ca^2+^ cations and stabilises unstable Mg-Ca calcite upon incorporating Mg^2+^ ions into the lattice.^109^ Calcium-binding IDPs have been classified into two groups according to their interaction motif: Excalibur-like or condensed-charge.^110^ Being promoted by negatively charged repetitions, the latter mechanism is assumed to engage with Ca^2+^ ions without compaction of the protein structure. Whereas Aspein unambiguously belongs to the condensed-charge category, interactions with Ca^2+^ ions instead trigger its chain collapse accompanied with the formation of turns. In that respect, the electrostatic collapse specifically observed with divalent metallic cations, mostly Ca^2+^-driven, strongly echoes the findings of Klepka B. P., *et al*.^111^ By combining biophysics with coarse-grained MD, they demonstrated that the structural ensemble of the aspartic and glutamic acid-rich protein (AGARP), an acidic IDOMP secreted by the scleractinian coral *Acropora millepora* (*A*. *millepora*), was shifted towards collapsed but unfolded conformers. Their compaction degree increased with cation valency, showing that, in addition to charge screening and neutralisation effects, Ca^2+^ and Mg^2+^ engage into specific chelating motif with the polyanionic IDP by bridging carboxylate moieties. The comparison of our system properties with those studied by Klepka B. P., *et al*. is all the more relevant, as AGARP and Aspein are both unusually acidic IDOMPs involved in the biomineralisation of marine lifeforms. While, coherently with Klepka B. P., *et al*., we did not observe the formation of long-range secondary structures, ions binding still induced local conformational constraints triggered by the formation of short-ranged and Ca-bridged turns along the SG(D)_n_ domain. To some extent, the interfacial cement protein MrCP20 from the *Megabalanus rosa* barnacle was shown, by coupling MD simulations with electron microscopy, to undergo similar and subtle structure changes allowing the sequestration of free Ca^2+^ and CO_3_^2−^ ions.^112^

However, the similarities with AGARP from *A. millepora* do not stop here, as Klepka B. P., *et al*. also proposed a LLPS biomineralisation pathway that strongly resonates with our MD data.^45^ Mechanistically, Aspein-driven CaCO_3_ condensation can be described as phase separation coupled to percolation **(Figure 10)**.^113^ First, Aspein molecules bind to Ca^2+^ and subsequently recruit CO_3_^2–^ ions into PPCC, Ca^2+^ cations acting as protein chain stickers (percolation). Through the mediation offered by Ca^2+^-carboxylate tethering with peripheral Asp, such assemblies coalesce to form condensates of higher sizes and density (phase separation). The resulting assembly initiates crystallisation by undergoing internal rearrangement that further dehydrates and clumps individually stabilised ACC precursors into calcite nuclei. Finally, calcite crystal growth is kinetically controlled by Asp-Ca^2+^ binding and the steady addition of PPCC to the dense protein-CaCO_3_ phase. In extrapallial conditions, whereas Aspein prevents the CaCO_3_ polymorphic switch induced by Mg^2+^ cations, the latter affect the lifetime of PPCC. Indeed, PPCC fusion is delayed upon addition of Mg^2+^ to the CaCO_3_ solution, hence favouring the persistence of PPCC or smaller condensates. Such effect is attributed to the occupancy of peripheral Asp sidechains by Mg^2+^ cations that reduces the fraction of surface-exposed carboxylate moieties available for PPCC coalescence, while cloistering such cations away from stabilised CaCO_3_ phases. Advocating for the stepwise association and phase separation of ions with IDOMPs during their biogenesis, Aspein is only expressed in the oyster mantle before being exported to the outer prismatic layer. In that respect, the Ca-bound proteins will encounter a carbonate-rich environment upon traversing the nacre region, where *P. fucata* carbonate anhydrases are primarily secreted for providing HCO^3−^/CO_3_^2−^ anions through the conversion of dissolved _CO2._114,115

As previously stated, such effects are mediated by the interplay between Aspein IDRs and its exceptional Asp density **(Figure 10)**. The more flexible DA and, to a lesser extent, Nter domains are assumed to act as diffusion barriers of Ca^2+^ ions thanks to their inherent plasticity and dynamical character, which regulates the addition kinetics of ions. As for the polyD region, the small DA domain was shown to remain unstructured in solution and to partly engage into ion binding. While being the primary Ca^2+^ binder, the SG(D)_n_ domain is also highly flexible before gaining in stiffness and compactness upon CO_3_^2–^ attraction and ACC stabilisation. For example, the DA IDR is suggested to play a step- and condition-wise dual role in the full-length protein: i) upon Aspein biogenesis, the DA domain is first fly-cast along with SG(D)_n_ to help catching and trapping Ca^2+^ cations; ii) DA serves as a diffusion barrier to control carbonate association with Ca^2+^ ions and their crystal nucleation within the collapsed SG(D)_n_ region.

Regarding the unique amino acid composition of Aspein, Asp has been reported to inhibit nucleation as well as serine residues, in a more efficient fashion than Glu amino acids.^116,117^ Furthermore, polyD additives have shown to have exacerbated effects on nucleation time with increasing chain length. Therefore, it is not surprising that Aspein, behaving like an exceptionally long polyD polymer, is an efficient inhibitor of CaCO_3_ nucleation by delaying CO_3_^2–^ incorporation and stabilising ACC phases. Additionally, Asp residues form a protective shell around ACC particles decreasing their dissolution probability in the classical dissolution-precipitation paradigm. As such, Aspein modulates ACC crystallisation pathway through a cooperative ion-association process. Further negative charges could be harboured by Ser residues through posttranslational phosphorylation, increasing the density of negative charges in the Ca^2+^-binding SG(D)_n_ domain. Phosphorylation was shown to be essential in the shell formation of *P*. *fucata*, as dephosphorylation of SMPs in the extrapallial fluid resulted in abnormal crystal growth and orientation within the shell layers.^118^

Aspein preferential affinity for Ca^2+^ over Mg^2+^ cations arguably arises from its exclusively aspartic content. Other studies have highlighted that Asp residues are stronger binders of Ca^2+^ and better fixators of biominerals compared to Glu by pulling more ions in their vicinity and fostering their clustering in a reduced space, especially for long polyD chains.^119–121^ Three main reasons have initially been proposed to explain such discrepancies that are hereafter extended to the context of CaCO_3_ biomineralisation: i) binding sites are distributed in a smaller volume around polyD than polyE domains, hence offering a higher ionic density for reaching supersaturation; ii) due to their shorter sidechain length, Asp residues display a smaller covering angle upon Ca^2+^ binding, facilitating complementary charge neutralisation by CO_3_^2−^ counterions to stabilise ACC; iii) a higher strain is exerted on the ring formed by two neighbouring Asp sidechains, resulting in the detachment of one carboxylate available for coordinating a second Ca^2+^ cation, which favours the creation of ion clusters for crystal nucleation. Owing to its polyelectrolytic IDP essence, the inherent flexibility and structural dynamics of Aspein also likely explain its selectivity for Ca^2+^ over Mg^2+^ cations, allowing some diversity in coordination number.^122^

Drawing from our results and previous reports, we underpin that Asp residues play a pivotal role in the polymorph selection of calcite from the extrapallial fluid of *P. fucata* and, to a greater extent, in other marine molluscs. Supporting such a claim, Prismalin-14, another acidic IDOMP of *P*. *fucata* devoid of Glu residues, is also secreted into the extrapallial space and specifically transported to the prismatic layer where it is responsible for CaCO_3_ nucleation thanks to its polyD domain, as well as chitin tethering.^123,124^ Comparatively, other disordered SMPs secreted in the nacreous layer of *P. fucata*, such as Pif80 or Pearlin, have a more balanced Asp/Glu ratio enhancing their ability to bind or incorporate Mg^2+^ cations and promoting the crystallisation of aragonite.^42,125,126^ Such tendencies have recently been acknowledged for Ca/Mg carbonates biomineralised by several marine bivalves.^127^ Coherently, the prismatic layer organic matrix is composed of 53 mol% of Asp, compared to the nacreous layer containing 45 mol% of Asp+Glu.^33^ Moreover, aragonite-selecting proteins from other clades, such as AGARP from the staghorn coral *A*. *millepora*, also display a higher content in Glu amino acids.^45^ To the best of our knowledge, this is the first time that a correlation between Asp/Glu compositional biases in acidic SMPs and their shell layer-specific secretion has been underlined for explaining their regulatory role in CaCO_3_ polymorph selection exerted through, amongst other effects, the preferential metal binding.

Apart from Aspein, other IDOMPs have been identified in the secretome of *P*. *fucata*. Illustratively, the N16 (or n16) protein is shown to be mainly intrinsically disordered in solution and to undergo disorder-to-order transitions towards conformers enriched in β-turns during CaCO_3_ mineralisation.^128^ In addition, they showed that N16 preferentially selects vaterite and aragonite crystals, which is in good agreement with the correlation pictured so far between the Asp/Glu content and shell layer secretion. While Aspein is an Asprich protein specifically precipitating calcite in the prismatic layer, N16 is a component of the nacreous aragonitic organic matrix that contains a balanced composition in Asp and Glu residues. However, the molecular mechanisms by which N16 mediates polymorph growth are still unexplored. Complementary with acid-rich proteins, polybasic polypeptides, such as the KRMP family unique to the *Pinctada* genus or Shematrins, also contribute to the formation of shell layers through CO_3_^2−^ fixation.^21,129^

Aspein might be an ancestral SMP retained in the shellome of the *Pinctada* genus during the evolutionary shift from a calcite (low Mg/Ca ratio) to an aragonite sea (high Mg/Ca ratio) that promotes the formation of the calcitic prismatic layer of marine bivalves in nowadays aragonite seawaters. Interestingly, the polyD domain of Aspein has been shown to be conserved across *Pinctada* species (*P*. *margaritifera*, *P*. *maxima*, and *P*. *radiata*) and pterioid homologs without correlated patterns, revealing that the density of Asp residues is more important for defining the biomineralisation function of the protein than their arrangement.^130,131^ Supporting the prevalence of charge density, our simulations with the SG(D)3 and SG(D)4 motifs showed that both motifs indiscriminately bind Ca^2+^ and stabilise CaCO_3_ clusters in PPCC with respect to the polyD stretch length. Finally, temperature elevation and ocean acidification, which echo current climate changes, exert adaptability pressure on pearl oyster populations by affecting the *aspein* gene expression and altering its shell ultrastructure.^132–136^ In that respect, investigating such parameters on Aspein structure and function, as well as other SMPs and IDOMPs, represents a great interest for understanding how *Pinctada* oysters and other mineralised marine organisms respond to the transformation of their environmental conditions.

## 5. Conclusions

Biomineralisation is a ubiquitous process necessary for the generation and repair of biologically active mineralised structures essential to sustain life. Evidence now more than ever suggests that LLPS-undergoing IDPs are critical modulators of biogenic minerals growth. Notwithstanding compelling *in vitro* studies, the structure-function relationships of such proteinaceous systems remain mechanistically elusive at the molecular level. In such context, the Aspein protein, secreted by the Japanese pearl oyster *P. fucata*, is an unusually acidic prismatic SMP that selectively precipitates calcite over aragonite in the Mg-enriched extrapallial fluid. Based on our sequence-based predictions and modelling, Aspein was revealed to be a fully IDP, existing as a polyanionic statistical coil, which prompted us to coin a novel distinction for SMPs, i.e. IDOMP, that integrates protein intrinsic disorder into the biomineralisation picture. All-atom MD simulations carried out in different saline environments highlighted ion-specific responses as well as how Aspein draws advantage from its Asp density and the structural fuzziness of its domains in the purpose of CaCO_3_ biomineralisation into calcite.

Mediated by structure-less collapse, bridged turns formation, high chain flexibility, and coordinated protein-ion phase separation, Aspein selectively binds Ca^2+^ cations, attracts CO_3_^2–^ anions, inhibits CaCO_3_ nucleation, sequesters CaCO_3_ clusters, stabilises and dehydrates ACC phases, as well as promotes calcite growth while keeping Mg^2+^ cations at its surface, therefore precluding ACC poisoning and aragonite formation. By fly-casting its binding polyAsp IDRs, Aspein offers a greater interaction surface for ions and enhances the encounter of Ca^2+^ and CO_3_^2–^ ions in a controlled fashion, thus enabling to reach CaCO_3_ supersaturation by locally increasing the concentration of such ionic species thanks to subsequent electrostatic collapse, chain percolation, and condensation. Asp sidechains exert multiple roles in Ca^2+^ catching, CaCO_3_ trapping, PPCC fusion, and Mg^2+^ cloistering that all participate in the kinetically controlled selection of calcite in current seawater that thermodynamically favours aragonite. Moreover, Aspein secretion restriction to the prismatic shell layer in the mantle outer edge correlates its exclusively aspartic sequence composition. Asp residues are more selective binders towards Ca^2+^ compared to Glu and are more prone to impair Mg^2+^ embedding into prenucleation clusters or calcite. Conversely, acidic IDOMPs secreted in the nacreous shell layer display a more balanced Asp-Glu proportion for specifically precipitating aragonite under the influence of Mg^2+^ cations.

Through the paradigmatic example of Aspein, we demonstrated that IDOMPs, thanks to their unique sequence biases and conformational plasticity, constitute important SMPs for modulating the selection and growth of CaCO_3_ polymorphs that organise the molluscan shells. More globally, we also underpinned the molecular principles and mechanisms governing protein-mediated CaCO_3_ biomineralisation in a context relevant to acidic IDOMPs and marine organisms. Beyond enhancing our understanding of such phenomenon at the molecular level, we also hope that our findings will resonate with soft matter approaches for controlling the synthesis of polymorphic crystalline materials via the development of IDOMP-inspired polymers. Finally, although MD simulations offer a promising platform for apprehending biological processes, much work is still needed to elucidate the dialogue between the protein secretome and biomineralisation processes in aquatic life, particularly involving IDOMPs of other bivalves, brachiopods, corals, urchins, cephalopods, or crustaceans all facing the rapid transformation of their physico-chemical environment.

## Data availability

All software, algorithms, and codes exploited in this manuscript are freely available online. Data will be made available from the corresponding author upon request.

## Author contributions

J.M. conceptualised and designed the computational study, set up the simulated systems, conducted all the predictions and molecular dynamics simulations, processed and analysed the data, as well as wrote the original draft of the manuscript. C.M. supervised the work in the capacity of laboratory director. A.M. helped in the data formal analysis. A.M., S.L., and C.M. validated the processed data, as well as reviewed and edited the manuscript. All authors have given approval to the final version of the manuscript.

## CRediT author statement

Julien Mignon: conceptualisation; formal analysis; investigation; methodology; visualisation; writing – original draft.

Antonio Monari: formal analysis; validation; writing – review & editing. Sonia Longhi: validation; writing – review & editing.

Catherine Michaux: funding acquisition; resources; supervision; validation; writing – review & editing.

## Acknowledgements

The authors are grateful to the PTCI high-performance computing resource of the University of Namur (UNamur). The present research benefited from computational resources provided by the Consortium des Équipements de Calcul Intensif (CÉCI), funded by the Belgian National Fund for Scientific Research (F.R.S.-FNRS) under grant n°2.5020.11 and by the Walloon Region, and made available on Lucia, the Tier-1 supercomputer of the Walloon Region, infrastructure funded by the Walloon Region under the grant agreement n°1910247. A.M. thanks ANR and CGI for their financial support of the present work through Labex SEAM ANR 11 LABEX 086, ANR 11 IDEX 05 02. The support of the IdEx “Université Paris 2019” ANR-18-IDEX-0001 is also acknowledged. C.M. is thankful to the FNRS for her Senior Research Associate position.

## Abbreviations

ACC: amorphous calcium carbonate
Asprich: Asp-rich protein
CH: charge-hydropathy
CDF: cumulative distribution function
FES: free energy surface
HMR: hydrogen mass repartition
IDP: intrinsically disordered protein
IDOMP: intrinsically disordered organic matrix protein
IDR: intrinsically disordered region
LLPS: liquid-liquid phase separation
MSD: mean square displacement
MD: molecular dynamics
PME: Particle Mesh Ewald
PPCC: percolated protein-calcium carbonate cluster
PBC: periodic boundary conditions
PC: principal component
PCA: principal component analysis
PDF: probability density function
pRDF: proximal radial distribution function
RDF: radial distribution function
Rg: radius of gyration
RMSD: root-mean square deviation
RMSF: root-mean square fluctuation
SMP: shell matrix protein
SASA: solvent accessible surface area.

