## Supporting Information for "Molecular dynamics insights into biomineralisation mediated by acidic intrinsically disordered proteins: a case study of molluscan Aspein from the pearl oyster *Pinctada fucata*"

–

| | Aspein-D1 wild-type | | | SG(D)$_{3-4}$ peptides mixture [SG(D)$_3$:SG(D)$_4$)] | | |
|---|---|---|---|---|---|---|
| | 10 mM NaCl | 10 mM $CaCO_3$ | 10 mM $CaCO_3$ 50 mM $MgCl_2$ | 50 mM NaCl | 50 mM $CaCO_3$ | 50 mM $CaCO_3$ 250 mM $MgCl_2$ |
| System net charge | −54 | −54 | −54 | −140 | −140 | −140 |
| Box volume ($nm^3$) | 3723.9 | 3723.9 | 3723.9 | 1000.0 | 1000.0 | 1000.0 |
| Water molecules | 118768 | 118776 | 117784 | 29852 | 30020 | 28713 |
| Water atoms | 356304 | 356328 | 353352 | 89556 | 90060 | 86139 |
| $Na^+$ cations | 76 | / | / | 168 | / | / |
| $Ca^{2+}$ cations | / | 49 | 49 | / | 98 | 98 |
| $Mg^{2+}$ cations | / | / | 111 | / | / | 140 |
| $Cl^-$ anions | 22 | / | 222 | 28 | / | 280 |
| $CO_3^{2-}$ anions | / | 22 | 22 | / | 28 | 28 |
| Ionic content | 98 | 71 | 404 | 196 | 126 | 536 |
| Number of chains | 1 | 1 | 1 | 20:20 (40) | 20:20 (40) | 20:20 (40) |
| Atoms per chain | 1273 | 1273 | 1273 | 63:75 | 63:75 | 63:75 |
| Protein atoms | 1273 | 1273 | 1273 | 2760 | 2760 | 2760 |
| Total atoms | 357675 | 357672 | 355029 | 92512 | 92946 | 89435 |
| Minimisation (ps) | 2335 | 2071 | 2321 | 1460 | 884 | 2117 |
| Equilibration (ps) | 125 | 125 | 125 | 125 | 125 | 125 |
| Production (ns) | 3*1000 | 3*1000 | 3*1000 | 3*1000 | 3*1000 | 3*1000 |
| Simulation time | 18 μs | | | | | |

Table S1 – Summary of the molecular dynamics metrics, parameters, and durations for the six simulated protein systems, comprising i) the wild-type truncated version of Aspein (Aspein-D1) modelled in 10 mM NaCl, 10 mM $CaCO_3$, and 10 mM $CaCO_3$ with 50 mM $MgCl_2$ ionic environments; ii) the multicomponent peptide mixture of SG(D)$_3$ and SG(D)$_4$ motifs, twenty copies of each randomly dispersed for a total of forty chains, modelled in 50 mM NaCl, 50 mM $CaCO_3$, and 50 mM $CaCO_3$ with 250 mM $MgCl_2$ ionic environments.

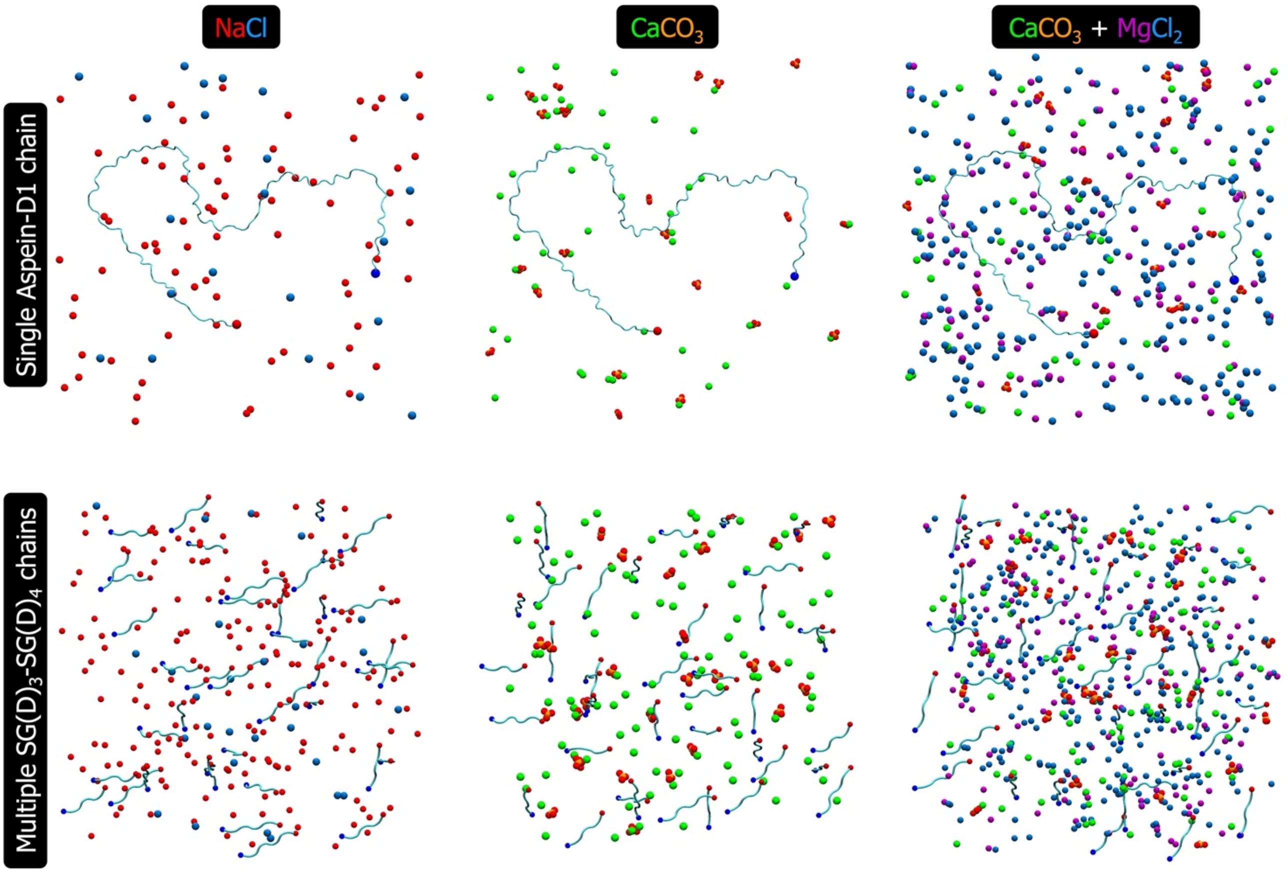


**Figure S1 – Initial states of Aspein single chain and multicomponent systems in different ionic environments**. Selected snapshots of Aspein-D1 wild-type (upper row) and SG(D)$_{3\text{-}4}$ peptides mixture (lower row) in tested extrapallial conditions, i.e. NaCl (left column), $CaCO_3$ (middle column), $CaCO_3$ with $MgCl_2$ (right column), after minimisation and equilibration. On the panels, $Na^+$ (red), $Cl^-$ (blue), $Ca^{2+}$ (green), $Mg^{2+}$ (purple), $CO_3^{2-}$ (orange and red for C and O atoms, respectively) ions are depicted as spheres, while the Aspein chains (cyan) are displayed in cartoon representation with the N- and C-termini respectively pinpointed as dark blue and dark red spheres.

A

[1]MKGIAILMCLAALVAVSVT[20]FPVADQTTNELGSSGAAAAGAVVSEPS[47]DAGDAADAGDADAADADAA
DADADADADADN[78]DDGDDDDDDDDDDSGDDDSGDDDDSGDDDDSGDDDDSGDDDDSGDDDDS
GDDDGDDDSEDDDDSGDDDSGDGDDGDSGDDDDDDDSGDDDDDDDSGDDDDDSGDDDDGDSG
DDDSGDDDGDDDDSGDDDSGDDDDSGDDDSGDDDDSGDDDSGDDESGDDDSGDDDDSGDDDS
GDDDSGDDGDDDDSGDDDDSGDDDDSGDDDDDDDDSGDDDDGDSGDDDSGDDDGDDDDSGDD
DSGDDESGDDDSEDDDSGDDDSGDDDSGDDDSDSGDDDSGDDDSGDDDSGDDDGDDGDDDADS
GDDDDDDDDDDGDDGDDDSGDDDGDDSDDDDDDDDDDQ[413]

Signal peptide N-ter DA SG(D)$_n$

B

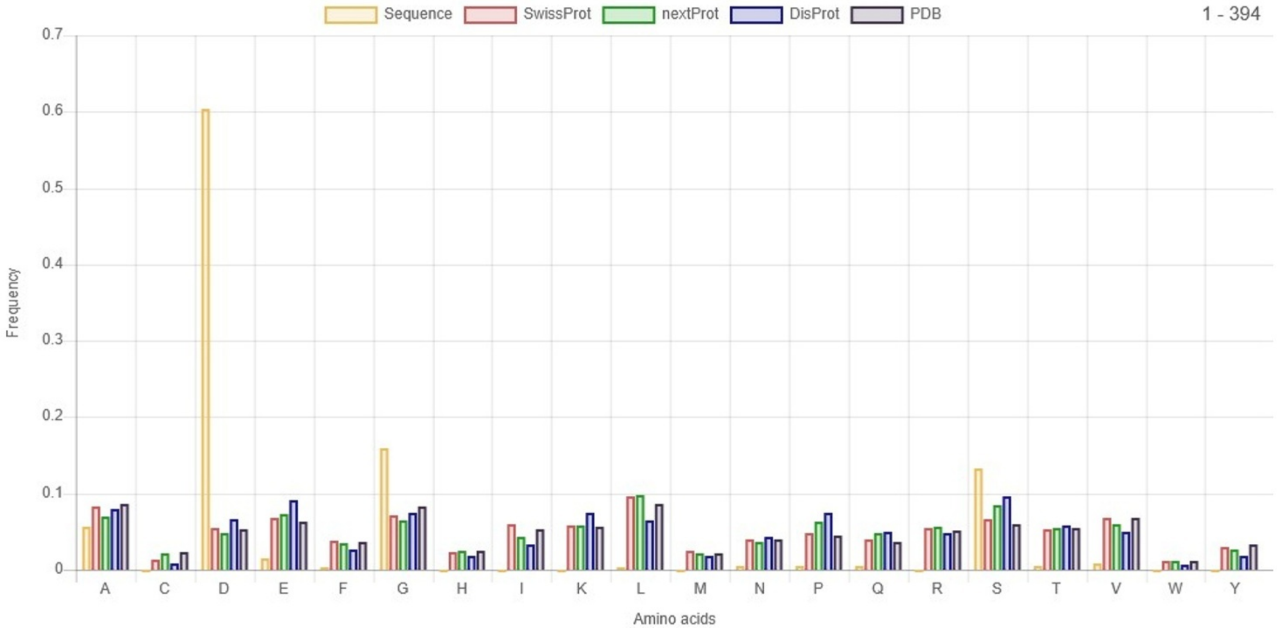


**Figure S2 – Sequence organisation of full-length wild-type Aspein from *P. fucata*.** (A) Per-residue sequence detail of immature Aspein with the cleaved signal peptide (green), as well as its functional Nter (grey), DA (orange), and SG(D)$_n$ (red) domains highlighted with respect to their position. (B) Relative amino acid abundance of mature Aspein (yellow) in comparison with the SwissProt (red), nextProt (green), DisProt (blue), and PDB (black) databases comprising ordered and disordered proteins.

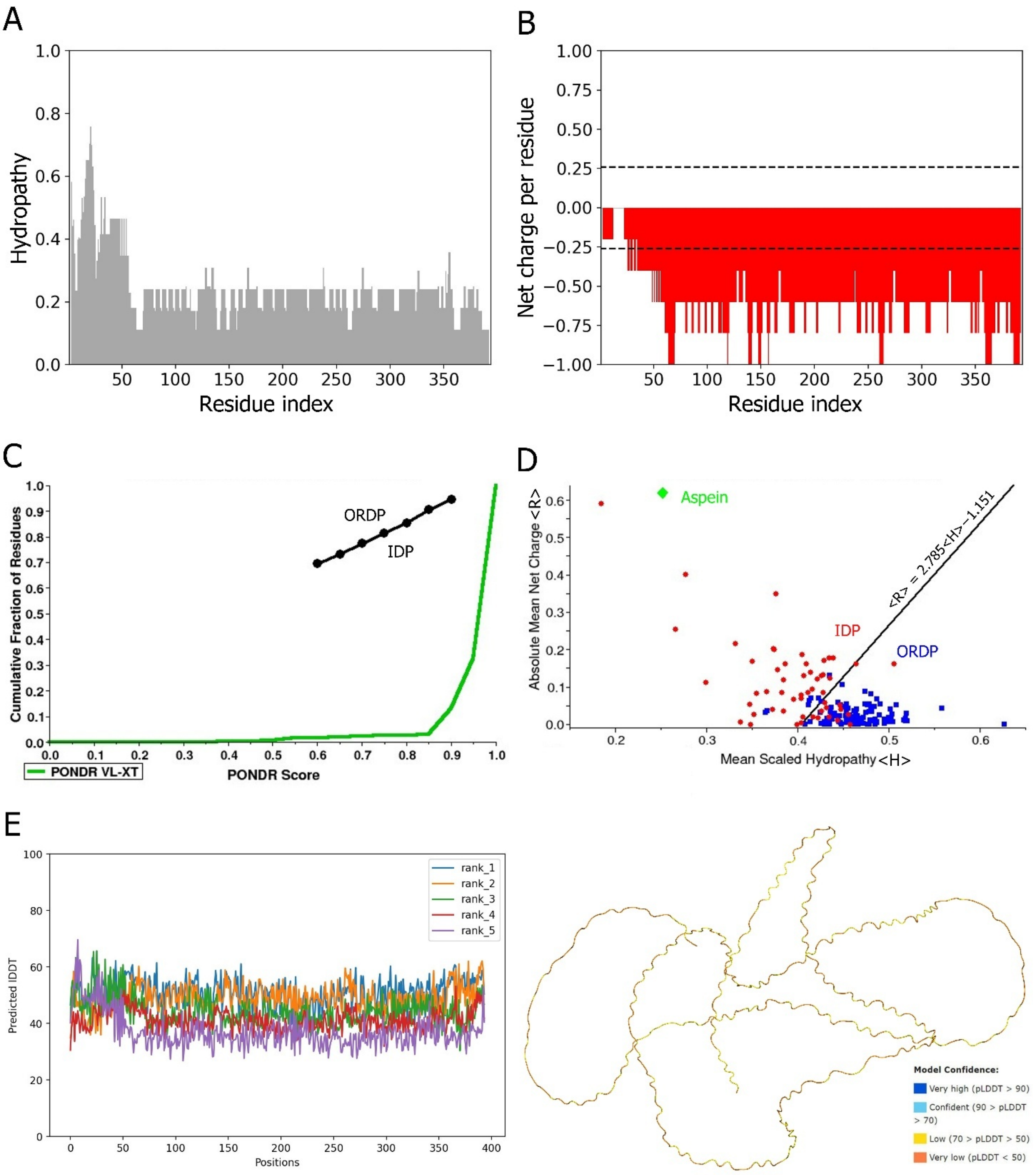


**Figure S3 – Hydropathy, charge distribution, and intrinsic disorder-associated predictions**. Linear distribution of (A) hydropathy (Kyte-Doolittle scale) and (B) net charge per residue using a sliding window of five residues. (C) Cumulation distribution function (CDF) and charge-hydropathy (CH) plots with the empiric boundaries between ordered (ORDP) and intrinsically disordered (IDP) proteins indicated. (E) MMseqs2-AlphaFold2 modelling of full-length wild-type Aspein structure showing the predicted confidence lDDT (scale at the bottom right) along the protein sequence (left) and projected on the rank 2 model displayed in cartoon representation (right).

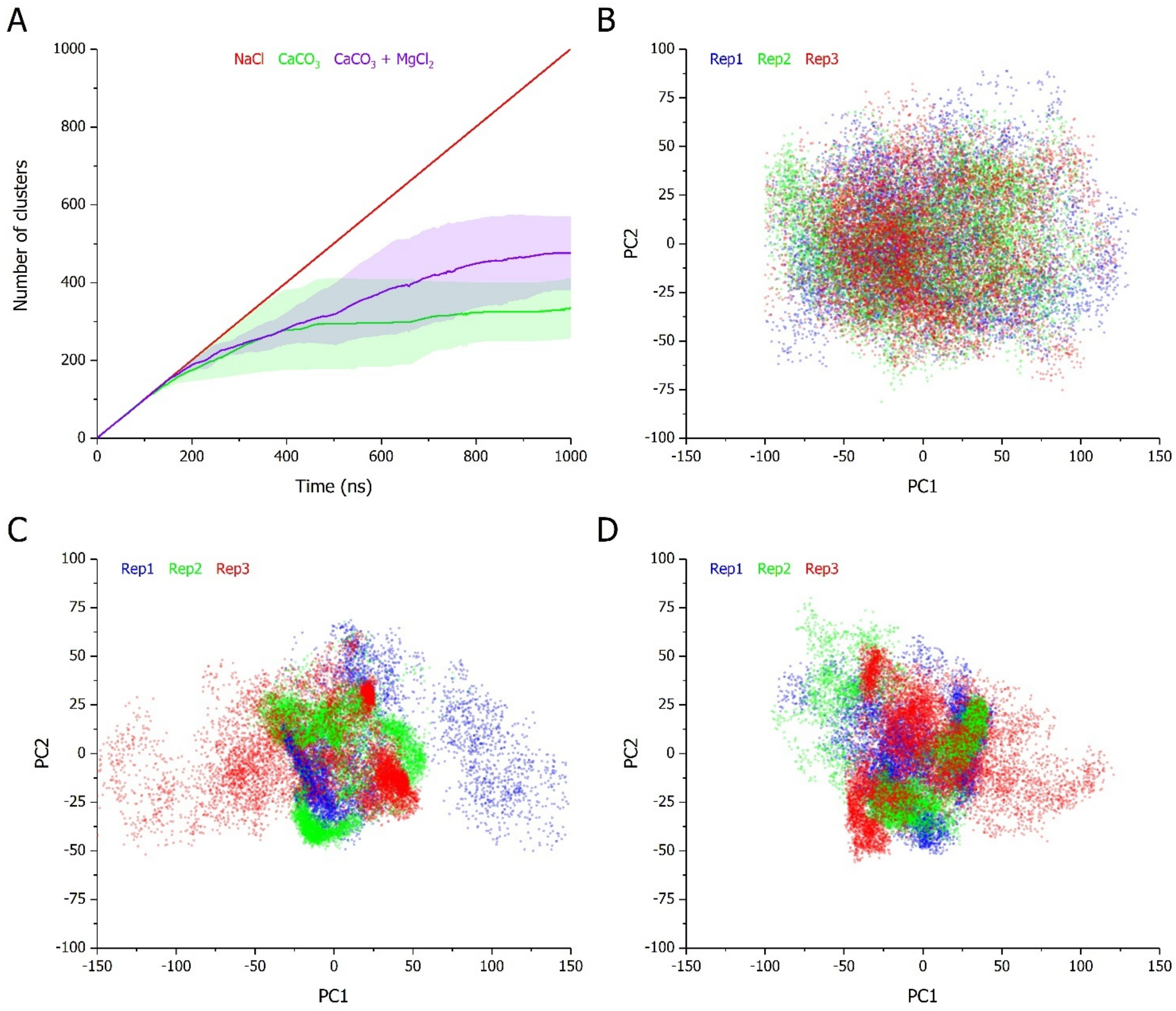


**Figure S4 – Clustering and PCA profiles of Aspein-D1 in different ionic environments**. (A) Time evolution of the number clusters using a 5 Å RMSD cut-off between the nearest structural neighbours of Aspein-D1 simulated for 1 µs in 10 mM NaCl (red), 10 mM $CaCO_3$ (green), and 10 mM $CaCO_3$ with 50 mM $MgCl_2$ (purple). On the panel, curves correspond to the average of triplicates with the standard deviation represented as a trace (shaded area) in the condition-associated colour. First two principal components (PC1, PC2) projection of Aspein-D1 simulated for 1 µs in (B) 10 mM NaCl, (C) 10 mM $CaCO_3$, and (D) 10 mM $CaCO_3$ with 50 mM $MgCl_2$. Each one of the triplicates (Rep1, Rep2, Rep3) is superimposed and distinctly coloured (blue, green, red).

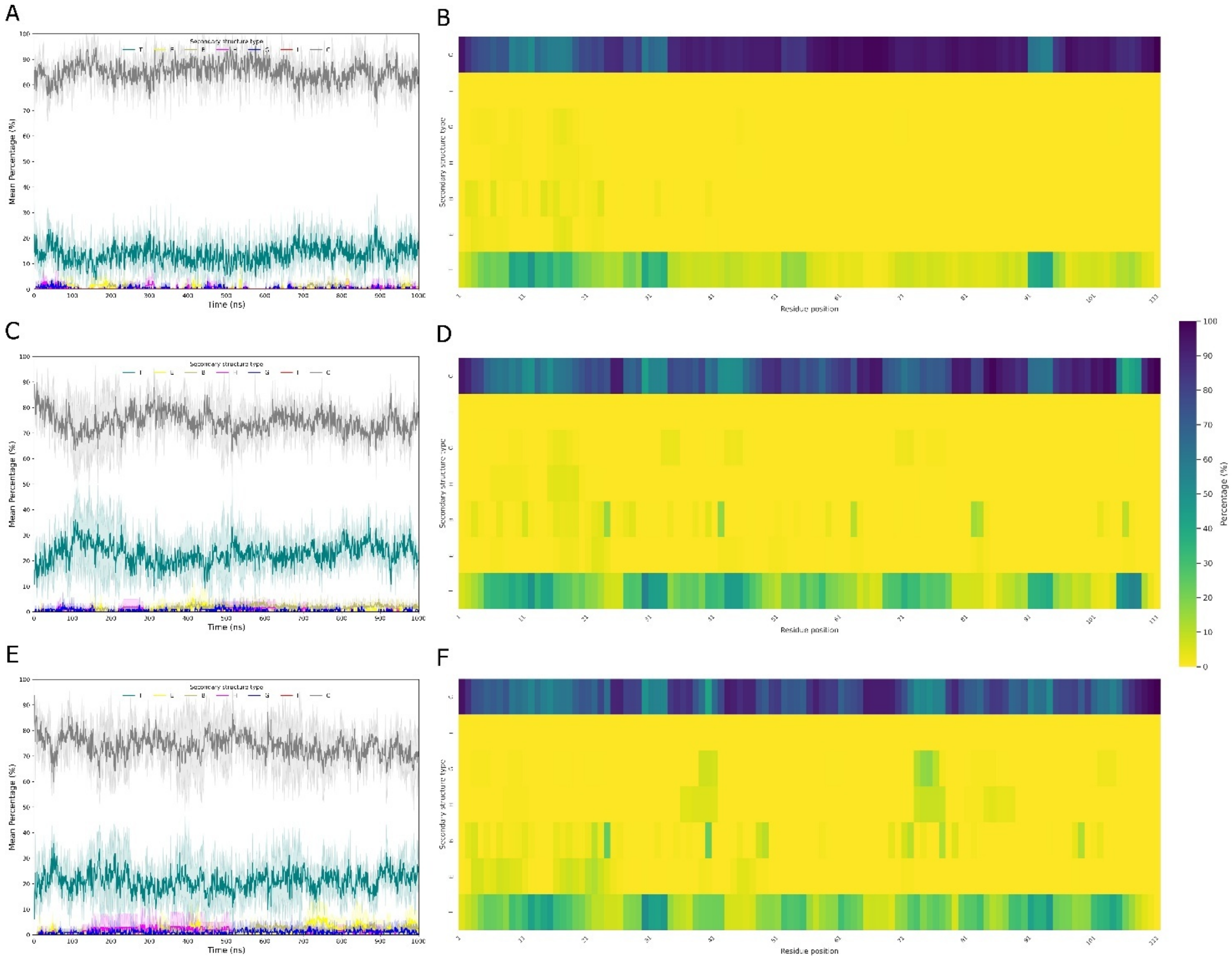


**Figure S5 – Diversity of secondary structures in Aspein-D1 in different ionic environments**. (A, C, E) Time evolution and (B, D, F) per-residue propensity of secondary structures of Aspein-D1 simulated for 1 μs in (A-B) 10 mM NaCl, (C-D) 10 mM $CaCO_3$, and (E-F) 10 mM $CaCO_3$ with 50 mM $MgCl_2$. The secondary structures are defined into seven classes after STRIDE assignment: turn (T), extended β-sheet (E), isolated β-bridge (B), $\alpha$-helix (H), $3_{10}$-helix (G), $\pi$-helix (I), and coil (C). On the time evolution panels, each secondary structure class is coloured as follows: T (green), E (yellow), B (dark yellow), H (magenta), G (blue), I (red), and C (grey); curves correspond to the average of triplicates with the standard deviation represented as a trace (shaded area) in the class-associated colour. On the occurrence panels, per-residue percentages correspond to the average of triplicates the scale of which goes from 0 (yellow) to 100% (dark blue) of propensity.

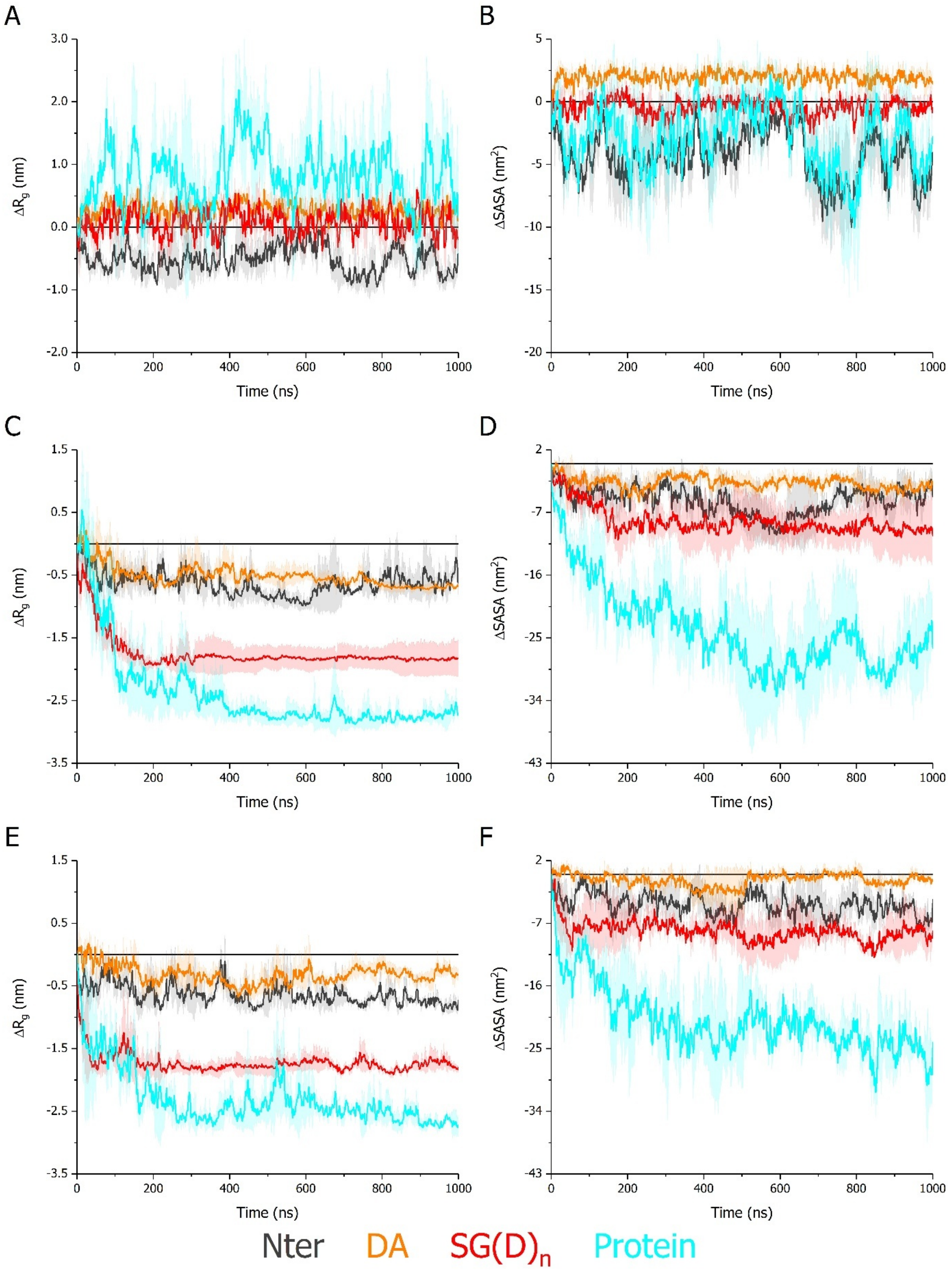


**Figure S6 – Compaction and hydration variation amongst the Aspein domains in different ionic environments**. Time evolution of (A, C, E) the radius of gyration ($\Delta R_g$) and solvent accessible surface area (ΔSASA) variation relative to the first timestep of the Nter (dark grey), DA (orange), and $SG(D)_n$ (red) domains, as well as the Aspein-D1 protein (cyan) simulated for 1 µs in (A-B) 10 mM NaCl, (C-D) 10 mM $CaCO_3$, and (E-F) 10 mM $CaCO_3$ with 50 mM $MgCl_2$. On each panel, curves correspond to the average of triplicates with the standard deviation represented as a trace (shaded area) in the domain-associated colour.

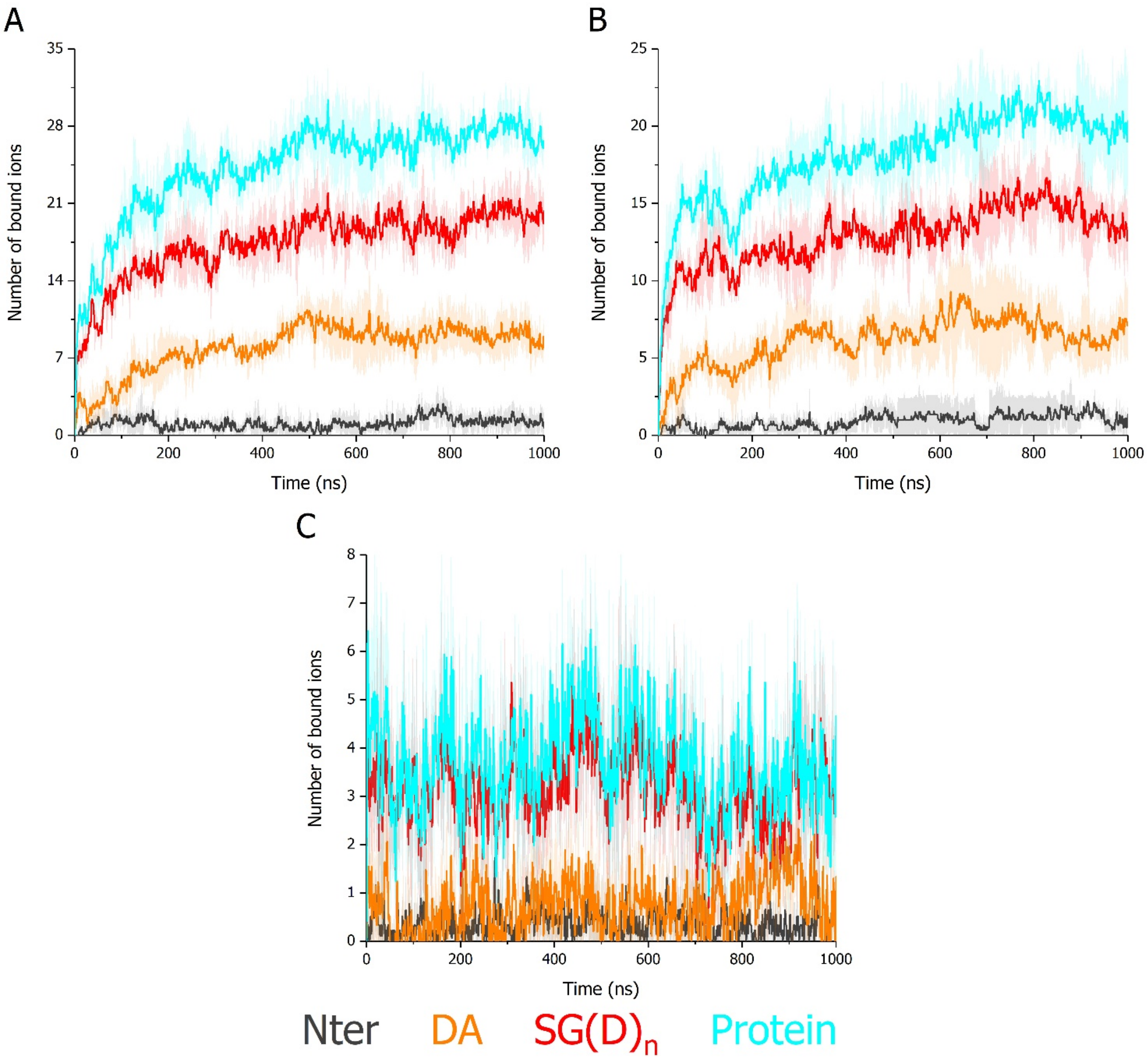


**Figure S7 – Selective ion binding amongst the Aspein domains in extrapallial conditions**. Time evolution of the number of contacts within a 4 Å cut-off around the protein surface involving the Nter (dark grey), DA (orange), and SG(D)$_n$ (red) domains, as well as the Aspein-D1 protein (cyan) with (A-B) $Ca^{2+}$ and (C) $Mg^{2+}$ cations encountered in the (A) 10 mM $CaCO_3$ and (B-C) 10 mM $CaCO_3$ with 50 mM $MgCl_2$ systems simulated for 1 µs. On each panel, curves correspond to the average of triplicates with the standard deviation represented as a trace (shaded area) in the domain-associated colour.

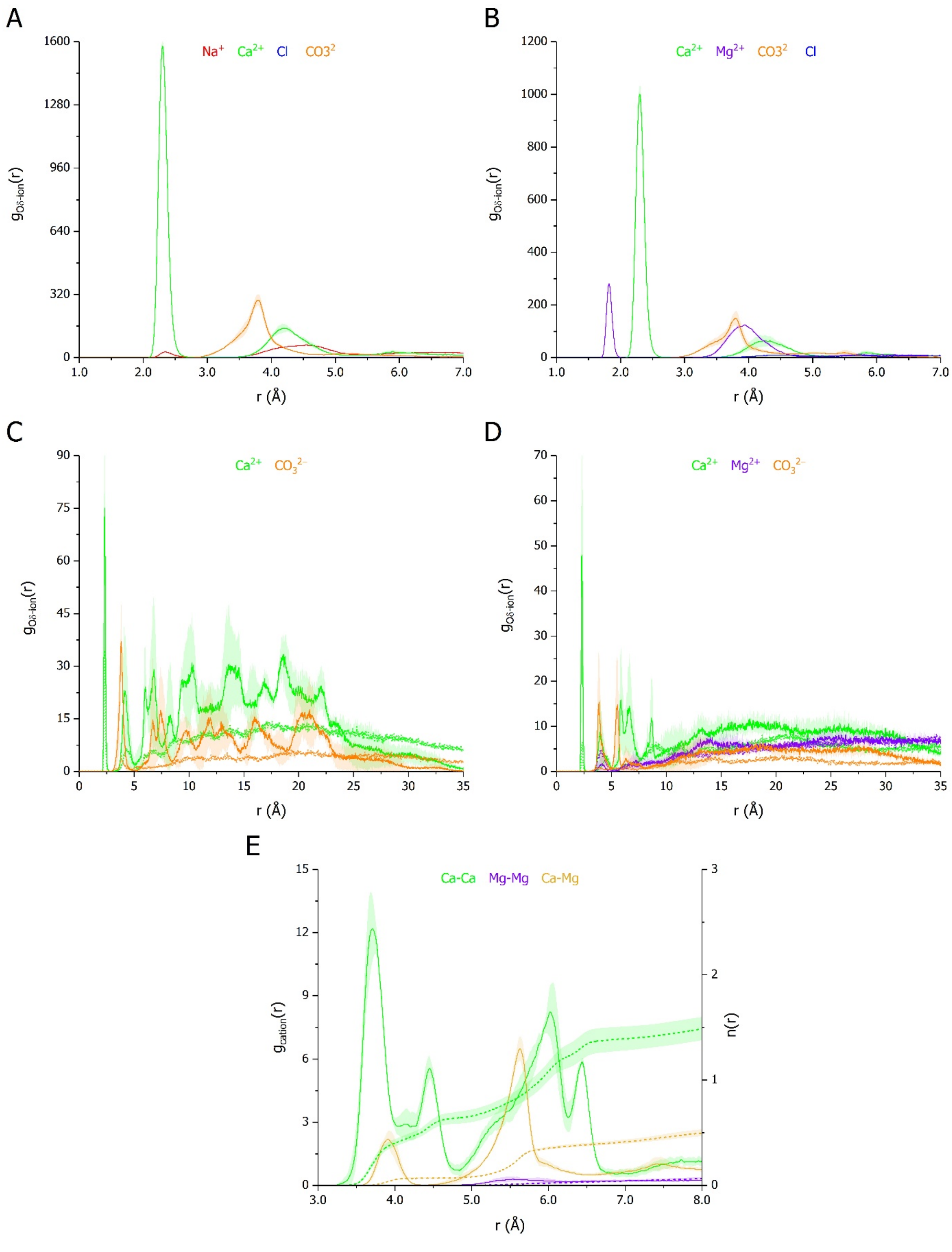


**Figure S8 – Ion distribution around Aspein Asp residues in different ionic environments**. Time-averaged proximal radial distribution function (pRDF) g(r) of $Na^+$ (red), $Ca^{2+}$ (green), $Mg^{2+}$ (purple), $Cl^-$ (blue), and $CO_3^{2-}$ (orange) ions with respect to (A-B) Oδ atoms of all Asp amino acids, (C) Oδ of D98, and (D) Oδ of D93 encountered in the Aspein-D1 systems simulated for 1 μs in (A) 10 mM NaCl, (A, C) 10 mM $CaCO_3$, and (B, D) 10 mM $CaCO_3$ with 50 mM $MgCl_2$. (E) Time-averaged pRDF (solid line) and cumulative number count n(r) (dashed line) of Ca-Ca (green), Mg-Mg (purple), and Ca-Mg (dark yellow) contacts for the Aspein-D1 simulations in 10 mM $CaCO_3$ with 50 mM $MgCl_2$. On each panel, curves correspond to the average of triplicates with the standard deviation represented as a trace (shaded area) in the ion-associated colour.

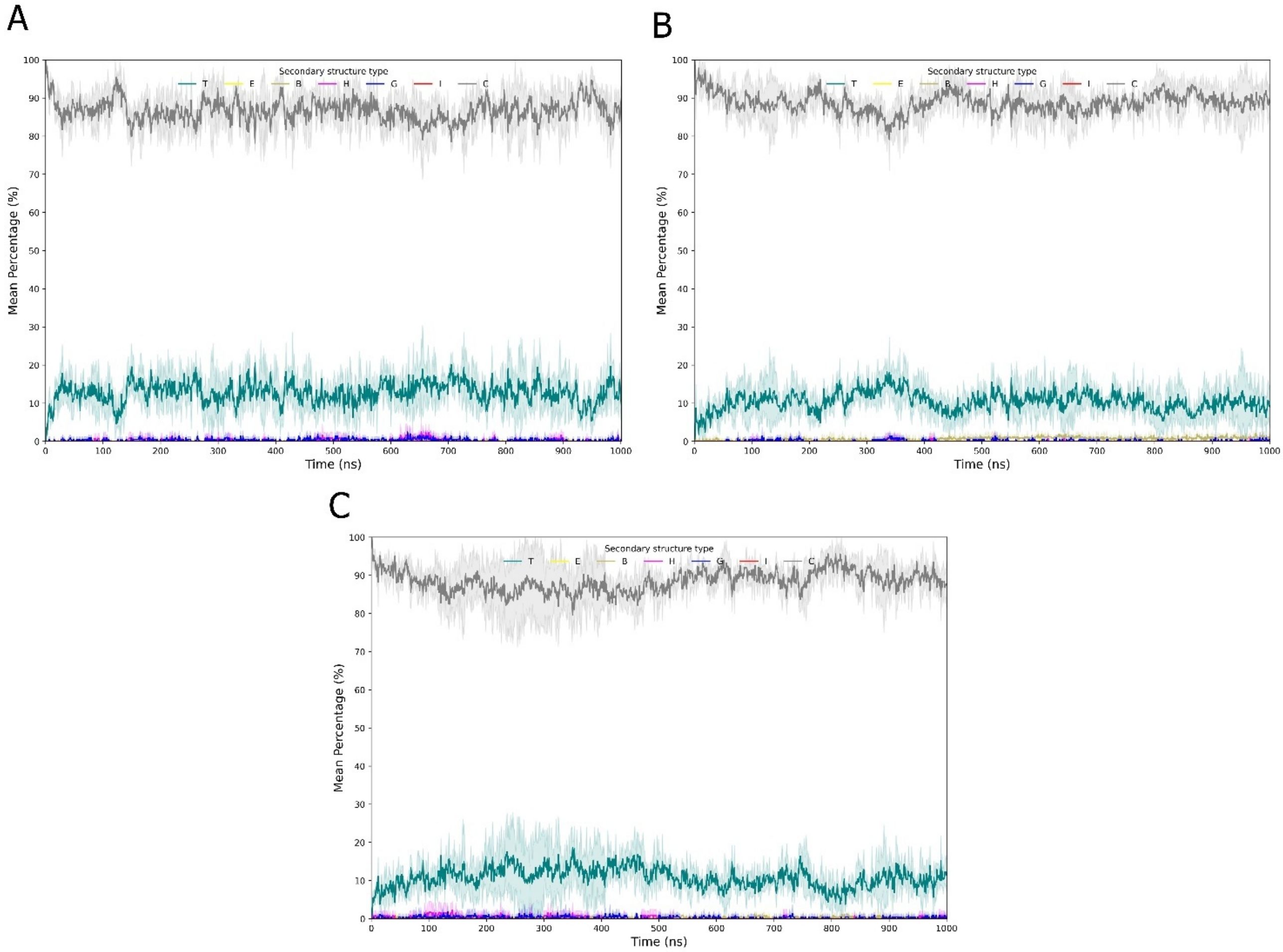


**Figure S9 – Diversity of secondary structures in the SG(D)$_{3-4}$ peptides mixture in different ionic environments**. Time evolution of secondary structures of the mixed SG(D)$_3$ and SG(D)$_4$ free monomers (twenty copies each) systems simulated for 1 μs in (A) 50 mM NaCl, (B) 50 mM $CaCO_3$, and (C) 50 mM $CaCO_3$ with 250 mM $MgCl_2$. The secondary structures are defined into seven classes after STRIDE assignment: turn (T, green), extended β-sheet (E, yellow), isolated β-bridge (B, dark yellow), $\alpha$-helix (H, magenta), $3_{10}$-helix (G, blue), π-helix (I, red), and coil (C, grey). On each panel, curves correspond to the average of triplicates with the standard deviation represented as a trace (shaded area) in the class-associated colour.

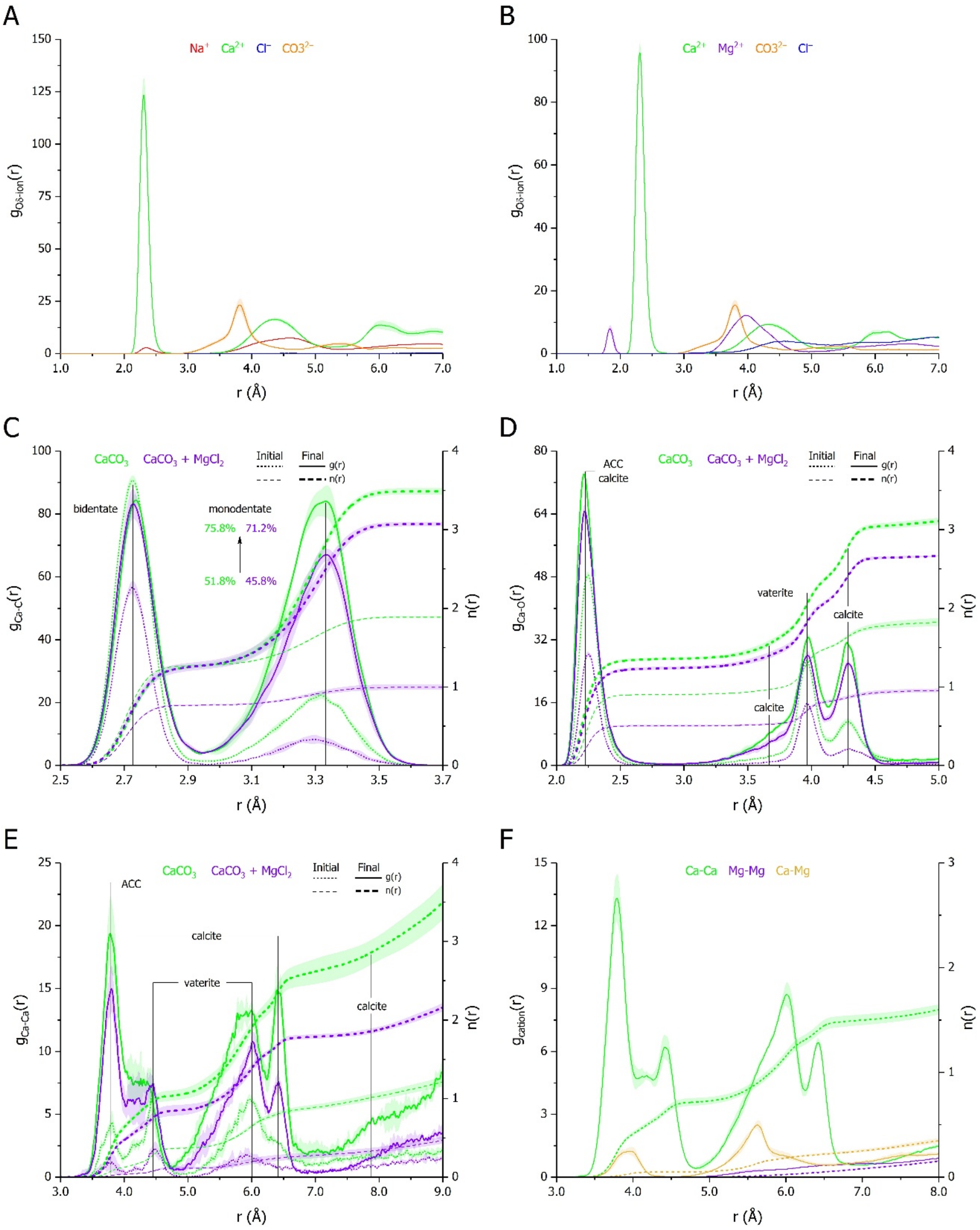


**Figure S10 – Selective effects of the SG(D)$_{3\text{-}4}$ peptides mixture on ion distribution and $CaCO_3$ polymorphism**. (A-B) Time-averaged proximal radial distribution function (pRDF) g(r) of $Na^+$ (red), $Ca^{2+}$ (green), $Mg^{2+}$ (purple), $Cl^-$ (blue), and $CO_3^{2-}$ (orange) ions with respect to Oδ atoms of all Asp amino acids encountered in the SG(D)$_{3\text{-}4}$ peptides mixture simulated for 1 μs in (A) 50 mM NaCl, 50 mM $CaCO_3$, and (B) 50 mM $CaCO_3$ with 250 mM $MgCl_2$. (C-E) Initial and final pRDF (dotted and solid lines) and cumulative number count n(r) (short-dashed and dashed lines) of (C) Ca-C, (D) Ca-O, and (E) Ca-Ca atoms from $Ca^{2+}$ and $CO_3^{2-}$ ions encountered in the 50 mM $CaCO_3$ (green) and 50 mM $CaCO_3$ with 250 mM $MgCl_2$ (purple) systems. (F) Time-averaged pRDF (solid line) and n(r) (dashed line) of Ca-Ca (green), Mg-Mg (purple), and Ca-Mg (dark yellow) contacts in 50 mM $CaCO_3$ with 250 mM $MgCl_2$. On each panel, curves correspond to the average of triplicates with the standard deviation represented as a trace (shaded area) in the ion-associated colour.
